# Evolution of temperature preference behaviour among drosophilids

**DOI:** 10.1101/2024.03.15.585210

**Authors:** Tane Kafle, Manuel Grub, Panagiotis Sakagiannis, Martin Paul Nawrot, Roman Arguello

**Affiliations:** Department of Ecology and Evolution, Faculty of Biology and Medicine, University of Lausanne, Switzerland; Computational Systems Neuroscience, Institute of Zoology, University of Cologne, Germany; School of Biological and Behavioural Sciences, Queen Mary University of London, London, UK

## Abstract

Small-bodied ectotherms are acutely vulnerable to temperature changes, but diverse thermotactic behaviours have contributed to their ability to inhabit broad climatic niches. Understanding how - and how quickly - these behaviours evolve are outstanding biological questions that are also relevant to conservation. Among insects, *Drosophila melanogaster* is a preeminent ectothermic model for temperate sensing and thermotaxis. However, little is known about how its temperature-related behaviours have evolved in comparison to its closely related species. We have thermo-profiled over 2400 larvae from eight closely related species of *Drosophila* from different thermal habitats. Consistent with local adaptation, we found substantial variation in temperature preference and fine-scale navigational behaviours amongst these species. Agent-based modelling of the larval thermotaxis circuit suggests that it is the balance between cool and warm avoidance circuits, rather than changes in temperature sensitivity, that drive differences in temperature preference. Our findings highlight the recurrent evolution of temperature-related behaviours in an experimentally tractable cross-species system.

## Introduction

Underlying the capacities that animals have to inhabit environments that are as variable as searing deserts^1–3^ and freezing polar regions^4–6^ are strategies to cope with temperature fluctuations that can vary extensively over short (e.g., seconds) and long (e.g., annual) timespans. For endotherms, strategies such as vasodilation, vasoconstriction and the use of brown adipose tissue, have evolved to maintain body temperature and metabolic balance thereby safeguarding homeostasis^7,8^. Such strategies that buffer internal body temperature against ambient temperature are largely unavailable to poikilotherms, animals that primarily rely on behavioural strategies such as moving up or down temperature gradients (thermotaxis), stopping in sunlight (basking), maximizing exposure to sunlight (flanking), or burrowing to regulate their internal temperature^9^.

The ability to make rapid behavioural changes for thermoregulatory purposes is particularly crucial for small-bodied poikilotherms, a group that includes most insects, as their internal temperature can match that of the environment within seconds due to rapid heat exchange^10–12^. The global distribution of ectothermic insects is a testament to their abilities to adapt thermoregulatory behaviours to their local conditions. These capacities have garnered extensive lab- and field-based research into the behavioural variation that exists within and between species across thermal environments^13–16^. Given that many small insects have been found to survive only within a narrow viable temperature range, an understanding of how fast such behaviours evolve - or how constrained they are - is increasingly relevant in light of rapid climatic change^17^.

A substantial part of our understanding of temperature-related adaptations in insects comes from work on drosophilids. Drosophilids are found in most places outside the polar regions and they have long served as study subjects for research on local adaptation. Initial work focused on cytological data that dates to Dobzhansky’s classic studies linking chromosomal inversions to climatic clines^18,19^, and has maintained a strong current to this day^20–23^. The behavioural and physiological studies that followed likewise began to document the roles that variable microclimates have in shaping diverse drosophilid species’ daily behaviours (e.g., hours of activity) and geographic distributions (with many species having very restricted ranges)^24–30^. Field observations, combined with collections that could be lab-maintained, helped to guide numerous thermotolerance experiments in which species’ ability to survive (or recover from) acute experimental temperature regimes could be readily assayed. This large body of work demonstrated remarkable differences among species’ abilities to survive both cold and hot temperatures and showed that these differences largely correspond with the thermal environments in which they are found^25,26,31^. Simple thermal gradient arenas and programmable Peltier elements have become increasingly common tools for quantifying temperature preference behaviours, principally in adult flies^32–34^. As with the tolerance experiments, these behavioural studies have identified large interspecies differences. For example, adults from a North American desert species, *D. mojavensis*, were found to prefer 27.9°C^28^ while adults from a high European alpine species, *D. nigrosparsa*, were found to prefer 10.4°C^35^. Adult *D. melanogaster* and *D. simulans*, two globally distributed ecological generalist species, prefer 24.3°C and 23.0°C, respectively^28^.

Temperature-related behavioural responses rely on the peripheral detection of thermal differences in the environment and the processing of that information by the central brain^36^. *D. melanogaster* is a preeminent neurogenetic model for thermosensation, and the characterisation of neural circuits and thermoreceptor proteins that underlie these behaviours is becoming increasingly complete, particularly with respect to the periphery. In adults, innocuous cool temperatures are detected by sensory neurons located in the antenna’s arista and sacculus^37,38^, with innocuous warm temperatures detected by another set of neurons in the arista and in the central brain’s anterior cells^37,39^. The neuron populations involved in cold and hot nociception in the adults are yet to be defined. In larvae, each dorsal organ ganglion, located in the head, houses distinct neuron populations that differentially respond to innocuous temperature changes by detecting ambient cooling or warming^33,40^. Noxious hot and cold temperatures are detected by multiple different classes of dendritic cells along the body wall of larvae^41–43^. The thermosensors that have so far been identified within these temperature sensitive neurons come from diverse families of ion channels including transient receptor potential channels, ionotropic receptors, and gustatory receptors, as well as members of the rhodopsin family; these have been detailed in recent review papers ^44,45^.

Most of the species that are closely related to *D. melanogaster* have narrower or non-overlapping climatic ranges^31,46–48^. The thermoecological diversity among these species, together with the cellular and genetic understanding of thermotaxis provided by *D. melanogaster*, put in place a strong foundation for comparative approaches to understanding the evolution of temperature-related behaviours^44,45^. Previous studies that have compared thermotaxis between drosophilids have primarily used distantly related species, which may have overlooked recurrent temperature preference changes if they evolve rapidly and limit phylogenetically-informed inferences about the history of the changes^28,49,50^. The few studies that have compared closely related species have focused only on a small number of target species^51^. As a result, it remains unclear how often temperature preferences evolve between species on short timescales. In addition, as most of this work has been carried out on adults, little is known about temperature-related behavioural evolution at the larval stage. Given the small size of larvae and their limited mobility, it is likely that selective pressures on thermotaxis at this developmental stage are distinct from those experienced by adult flies.

To address these questions, we have carried out a large larval thermotaxis experiment using eight species from two sister subgroups within the *D. melanogaster* species group: the *D. melanogaster* subgroup (hereafter abbreviated *Dmel*-subgroup) and the Oriental subgroup (Fig 1A). We focused on these two subgroups due to the inclusion of, and evolutionary proximity to, *D. melanogaster* and due to the evidence that multiple species within these two clades are believed to have recently experienced lineage-specific temperature-related adaptations^46,52–54^. We aimed to investigate if there is evidence that behavioural adaptation accompanied these changes. Our balanced species sampling from these two subgroups, together with divergence times that span relatively short to intermediate ranges, provide a powerful framework to investigate the rate and repeatability of thermotaxis evolution.

**Figure 1:**
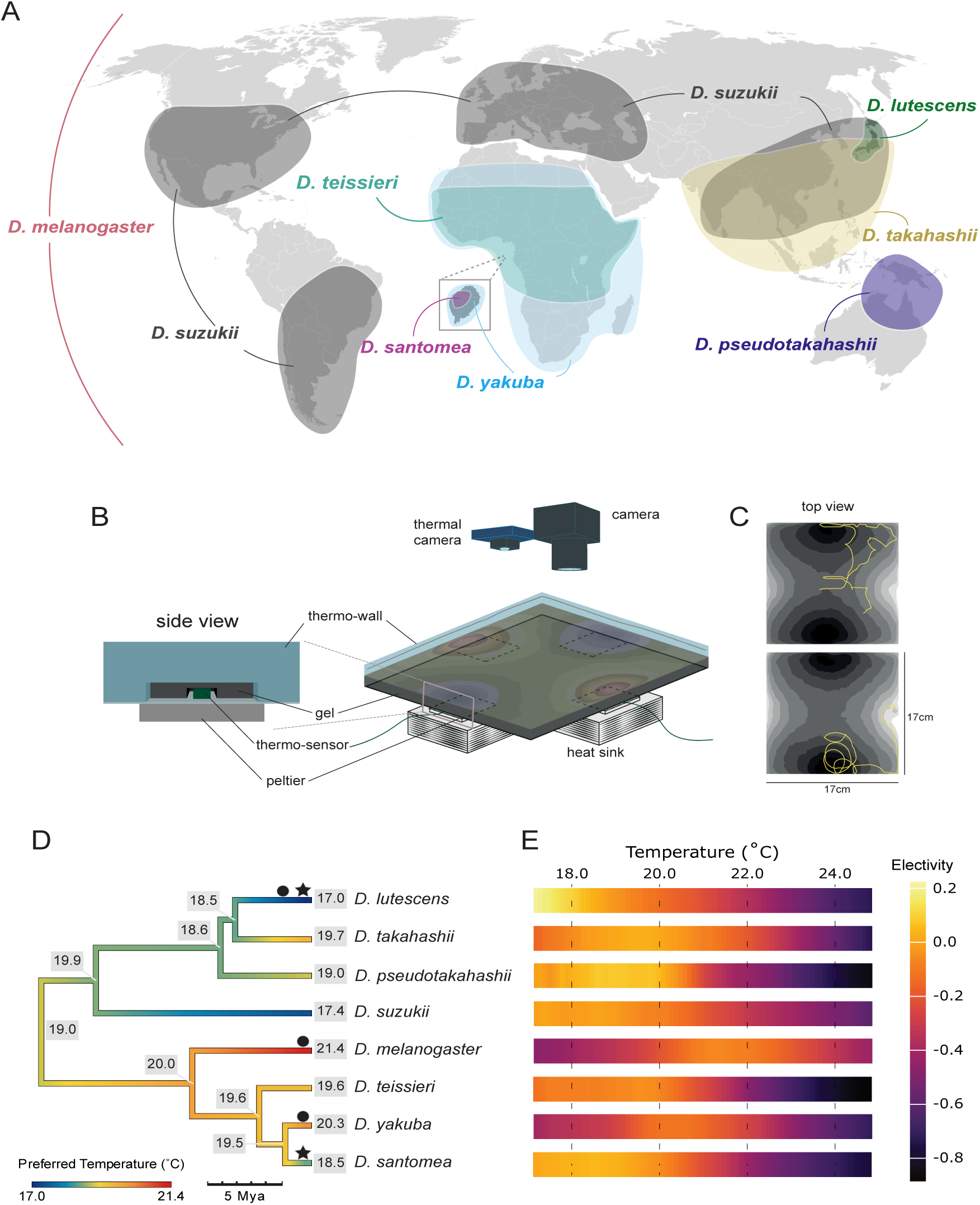
Ecologically diverse *Drosophila* species exhibit vastly different behaviours in relation to temperature. (A) Estimated ranges of the species used in this study. *Drosophila melanogaster* is found across the globe^59^, while the invasive pest species *D. suzukii* is found on most continents and is currently undergoing a global range expansion^60^. *Drosophila yakuba* and *D. teissieri* overlap for most their ranges on the African continent, with the former extending further into southern regions of the continent^51^. *D. santomea* is endemic to the island of Sao Tome off the coast of west Africa^48^. *Drosophila pseudotakahashii* is found primarily in northern Australia, and *D. lutescens* is found mainly in Japan^61^. *Drosophila takahashii* is also found in Japan but has a much greater range extending across mainland Asia^61^. (B) Schematic of the behavioural arena. The left panel shows a magnification of the side of the gel and plate. A small temperature sensor is placed between the plate and the gel above each Peltier element. This feeds back temperature recordings to an Arduino device, which in turn alters power fed to the Peltier elements to control the temperature. The right panel shows the arena with a thermal gradient overlaid on top of a black gel on an aluminium plate. Underneath the plate are four Peltier elements which are the temperature sources for the arena. Two cameras were placed above the arena, one to record larval movement and one to record the surface temperature. (C) Example tracks from two different larvae on the temperature gradient shown in panel B. Lighter colours represent warmer temperatures and darker colours represent cooler temperatures. The first track shows a weak Electivity to the darker regions of the gradient (i.e. weak preference to cool temperatures), whereas the second track shows a very strong Electivity to the darker regions (i.e. strong preference to cool temperatures). (D) Dated phylogeny of the eight species used in this study coloured by *E_peak_* values. Black stars indicate a shift found in *E_peak_* and black circles demonstrate shifts found in the upper limit of *E_breadth_*. A reduction in *E_peak_* values for both *D. lutescens* and *D. santomea*, indicates that they prefer cooler temperatures than their closely related species. Additionally, a downward shift of the upper limit of *E_breadth_* in *D. lutescens* indicates that it displays warm avoidance behaviours at lower temperatures than other species. In contrast, *D. melanogaster* and *D. yakuba* have increased their upper limits of *E_breadth_*, indicating they spend more time at warmer temperatures than their sister species. Ancestral *E_peak_* values are displayed at the nodes of the species tree (Methods). (E) Heatmap for each species showing their Electivity scores across temperatures tested in our experiments with lighter (orange, yellow) shades representing positive Electivities (preference) and darker (purple) shades representing negative Electivities (avoidance).

To quantify temperature-related behaviours in larvae, we implemented a novel temperature gradient assay paired with high resolution individual tracking. These data allowed us to continuously monitor broad patterns of species’ thermotaxis, as well as individual’s fine-scale behaviours throughout each experiment, details that have previously only been collected for *D. melanogaster*^55,33^. Analysing these data within a phylogenetic context, we have identified recurrent evolutionary changes within both species subgroups. Fitting species-specific models of larva thermotaxis to our data, we found evidence that evolutionary changes between species are explained by differences in the balancing of signals from the cool and warm circuits in the larval brain and not by changes in sensitivity to cooling/warming temperatures.

## Results & Discussion

### Recurrent changes in temperature preference

Thermal environment at the microhabitat scale (the scale of a single plant or fruit) is an important behavioural determinant of small ectotherms^56,57^. To provide a realistic “thermoscape”, similar to what is experienced by crawling insects in the wild (e.g., moving into/out of sun/shade over short distances), we developed an assay that tracks the movements of individuals within a 17 x 17cm thermal arena that was designed to hold patchy non-noxious temperature gradients on its surface^56^ (Figure 1B-C). We collected temperature-profiled tracks for third-instar larvae from eight species: *D. lutescens*, *D. takahashii*, *D. pseudotakahashii*, *D. suzukii*, *D. santomea*, *D. yakuba*, *D. teissieri* and *D. melanogaster*. Three strains were used for each species, except for *D. pseudotakahashii*, for which we could only obtain a single strain. In total, we collected 3884 larvae, assayed across 191 independent 20-minute experiments (Methods; Table S1). Following quality control filtering that, among other criteria, ensured that all gradients maintained a temperature range of 17-25°C, a dataset of 2418 larva tracks from 129 experiments remained for analysis. Each species’ temperature preference was quantified using Ivlev’s Electivity^58^ (Methods, Equation 1). Ivlev’s Electivity (*E*) is a common preference index used in the foraging literature that accounts for uneven resources and is well-suited for our analysis due to the unequal temperature bins generated over the surface of our arena. *E* ranges from -1 to 1, where -1 denotes a strong avoidance of those temperatures, and 1 denotes a strong preference for those temperatures. Because temperature bins that are never (or rarely) explored result in negative *E* values (and because larvae can only be within one temperature bin at a given time) our estimates of temperature preference tend to be negatively biased (see Figure S1 for additional details).

Our initial examination of temperature preferences over the eight species revealed significant variation between species in peak preference (*E_peak_*) and the breadth of the preferred temperature range (*E_breadth_*). Temperature preference varied significantly more between species than within species, indicating that substantial genetic change has occurred over the diversification of the eight species for this trait (ANOVA on *E_peak_* : *F* (6,18) = 7.43, *p*-value <0.01; Fig. S2-3, Table S2). Despite the negative bias for *E* in our experiments, all Oriental clade species, along with *D. santomea*, had significantly positive mean Electivity values at *E_peak_*, indicating strong preference for those temperatures (Table 1). Interestingly, in each case the strong preference was for the cooler temperatures available in the arena (Fig. 1D). In contrast to the aforementioned six species, *D. melanogaster* and *D. yakuba* had relatively low values of Electivity at *E_peak_* which occurred at warmer temperatures, indicating that they have a comparably weak temperature preference over the range tested (Table 1).

**Table 1:** Electivity measures across species. *E*_peak_ is defined as the temperature (°C) with the highest mean Electivity. The Mean column provides the mean Electivity value of *E*_peak_ for each species. Values in the *E*_breadth_ column provide the range of temperatures where larvae spent time, calculated using a sign test on electivity values comparing larval movement across temperatures to random movement (Methods). The *E*_breadth_ (strong) column shows the temperature range where larvae spent significantly more time than expected based on a from the sign test (Methods). NA indicates that there was no temperature range in which the species showed a strong preference.

| Species | $E_{peak}$ (°C) | Mean ($E$) | $E_{breadth}$ (°C) | $E_{breadth}$ (strong; °C) |
| --- | --- | --- | --- | --- |
| *D. lutescens* | 17.0 | 0.226 | 17.0 - 19.3 | 17.0 - 18.6 |
| *D. takahashii* | 19.7 | 0.034 | 17.0 - 21.7 | 18.8 - 20.2 |
| *D. pseudotakahashii* | 19.0 | 0.084 | 17.0 - 20.9 | 19.8 |
| *D. suzukii* | 17.4 | 0.005 | 17.0 - 20.5 | 17.0 |
| *D. santomea* | 18.5 | 0.050 | 17.0 - 20.5 | 17.0 - 19.3 |
| *D. yakuba* | 20.3 | -0.113 | 19.4 - 22.1 | NA |
| *D. teissieri* | 19.6 | -0.047 | 17.0 - 20.6 | 17.7 - 17.8 |
| *D. melanogaster* | 21.4 | -0.065 | 20.2 - 23.3 | NA |

Among the most notable differences in temperature preferences are the prominent cool-preferences observed for *D. lutescens* (*E_peak_* = 17°C (lower limit of study), *E_breadth_* = 17.0-19.3°C) and *D. santomea* (*E_peak_* = 18.5°C, *E_breadth_* = 17.0-20.5°C). In both cases, the preferences are significantly different when compared to their sister species (*D. takahashii* and *D. yakuba*, respectively) indicating that the changes happened relatively recently: *D. lutescens* preference is significantly higher than *D. takahashii* between 17.0-17.8°C and *D. santomea* preference significantly higher than *D. yakuba* between 17.0-19.6°C (both MWU tests *p*<0.01). These species-specific preference changes for cooler temperatures are intriguing because *D. lutescens* and *D. santomea* are both found in cooler climates compared to their respective sister species^31,48^ and because past work has independently provided evidence that they have adapted to cooler climates. For example, even after exposing *D. lutescens* larvae to near freezing temperatures (3°C) for over a month, larvae are viable and able to eclose after pupating^30^. In contrast, *D. takahashii* larvae die under the same conditions within four days^30^. The increased cold temperature resistance in *D. lutescens* compared to *D. takahashii* is also observed in adults^31,54^. Our results suggest that behavioural changes have coevolved with physiological adaptations to enable *D. lutescens* to live in colder environments. Similarly, adult *D. santomea* prefer cooler temperatures compared to *D. yakuba* and each species suffers fitness costs if reared at the other’s preferred temperature (particularly *D. santomea*)^52,62^. Our findings expand upon these observations by demonstrating that temperature preference behaviour spans both adult and larval stages.

Beyond *D. lutescens’* and *D. santomea’s* cooler preferences, the broader variation that we observed in temperature Electivity suggested further changes in the history of the eight species (Fig. 1D). Between the two subgroups we found that the Oriental clade species have stronger preferences (higher Electivity) at cooler temperatures than species of the *Dmel*-subgroup (MWU tests *p*<0.001 for temperatures below 19°C). To test for changes in temperature preference more generally, we modelled the evolution of *E_peak_* and the onset of warm avoidance (the upper bound of *E_breadth_*) within a phylogenetic context and asked if there is evidence of significant changes in either metric along any of the branches in the species tree. Due to limitations of the arena’s design we were unable to carry out the same tests for the lower bound of *E*_breadth_. These analyses provided additional confirmation of the changes in *D. lutescens*’ and *D. santomea’s E*_peak_, with estimated lineage-specific shifts of - 2.34°C and -1.68°C, respectively, compared to their inferred ancestral values. For *D. lutescens*, this is particularly notable given the conservative estimate of its *E*_peak_. Intriguingly, we identified additional parallel preference shifts impacting warm avoidance (upper bound of *E*_breadth_) for *D. lutescens*, *D. melanogaster*, and *D. yakuba*. The onset of heat avoidance has evolved to be 1.87°C lower in comparison to the inferred ancestral value for *D. lutescens*, suggesting a decrease of the upper tolerance bound of its preference of ∼17°C. In contrast, the onset of heat avoidance has expanded for both *D. melanogaster* and *D. yakuba*, indicating that both spend significantly more time within warm zones in comparison to the estimate inferred for their respective common ancestors. We found the change in the upper *E*_breadth_ for *D. melanogaster* to be 1.56°C higher than the inferred ancestral value, while the same estimate was 2.66°C greater for *D. yakuba’s* (Fig. 1D). These results highlight the recurrent and fast rates at which the peak and breadth of temperature preferences have evolved among larvae of closely related *Drosophila* species.

### Navigational metrics support the recurrent evolution of temperature preference

In addition to the relative amount of time *Drosophila* larvae spend within temperature zones, fine-scale individual navigational behaviours are also reflective of thermal preference and avoidance. During positive taxis, larvae move with relatively direct linear motion in comparison to negative taxis during which they move more tortuously, reflecting attempts to stay within preferred temperatures^63^ (Fig. 2A). While agent-based modelling has shown that changes in the rate of turning can capture most larval taxis behaviour^64^, they also vary speed whilst navigating (Fig. 2B). For example, in response to olfactory cues, larvae modulate speed in response to aversive and attractive odour gradients^65^, and similar changes in speed have been observed on thermogradients^66^. Additionally, because a larva’s non-noxious thermosensors are located at the tip of their head^44^ (along with sensors that detect other environmental cues ^67,68^), the initiation of turns is established by first probing their environment using head sweeps. As negative thermal stimuli evoke larger turns^33^, larger head sweeps are expected to reflect increasingly aversive temperatures compared to head sweeps in preferred temperatures^33^ (or other favoured stimuli; wind^69^, light^70^, olfactory^71^) (Fig. 2K).

**Figure 2:**
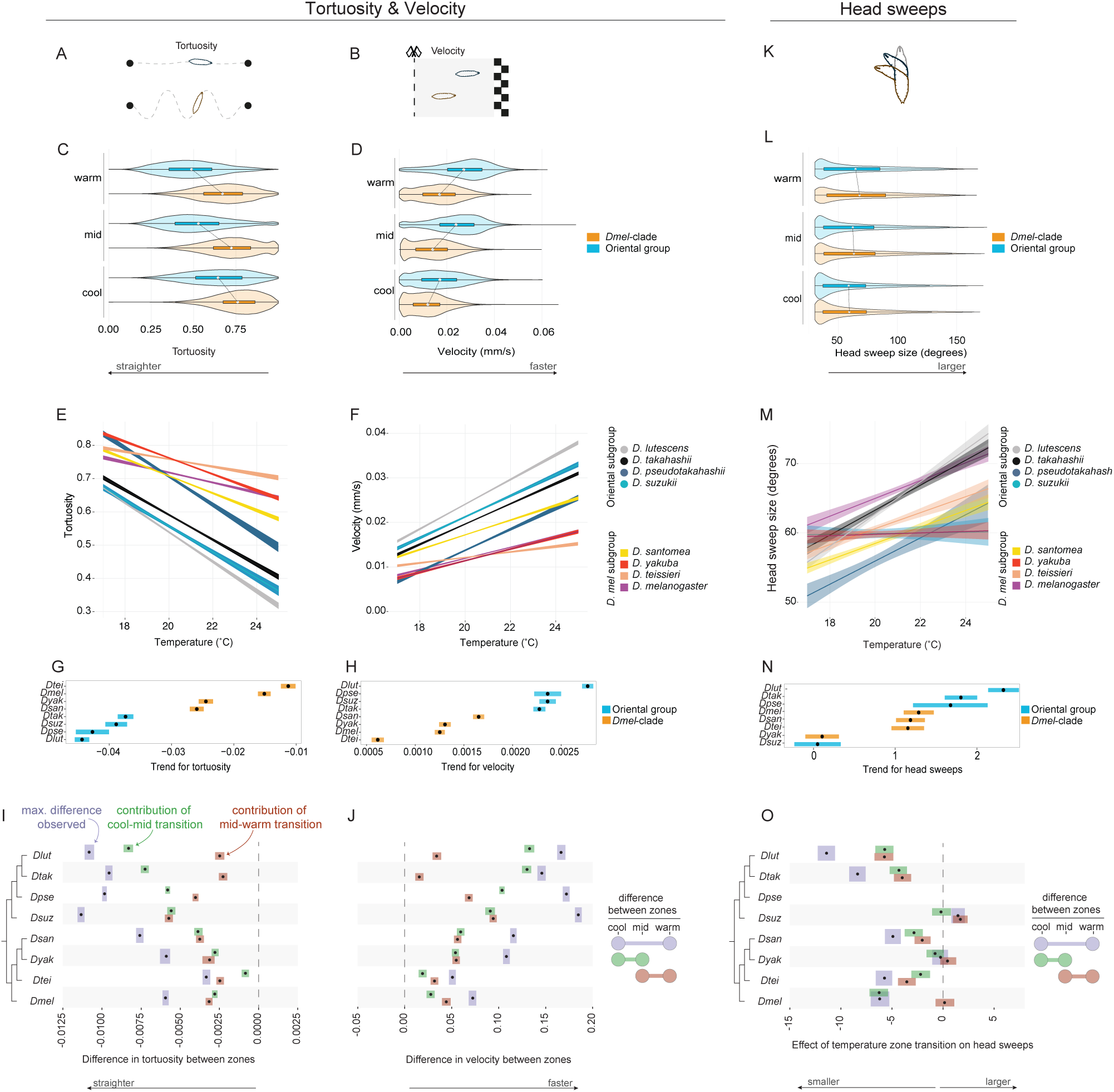
Fine scale navigational metrics across temperatures. (A,B,K) Illustrations of the navigational metrics that were measured for individual larvae: velocity, tortuosity and head sweeping. Velocity and tortuosity are highly correlated and therefore displayed adjacently. (C,D,L) Comparisons of navigational metrics between pooled *Dmel*-subgroup and pooled Oriental clade individuals across cool (17.00-19.67°C), mid (19.67-23.33°C), and warm (23.33-25.0°C) temperature zones. Violin plots are overlaid on top of box plot where the white circle represents the mean, and the hinges represent the first and third quartiles. Significant clade-differences are observed for velocity and tortuosity, particularly at the warm and mid temperatures. Differences among clades were not observed for head sweeps. (E,F,M) Linear fits of the behavioural data across the thermal gradient’s temperatures. Shaded envelope surrounding are lines are the 95% confidence interval. Tortuosity R^2^ = 0.279, Velocity R^2^ = 0.266, Head sweep size R^2^ = 0.0159. (G,H,N) Estimated means (black circles) and 95% confidence intervals (coloured bars) for the trend (slopes) of the linear fits shown in panels E, F, M. The trends separate the *Dmel*-subgroup from the Oriental group for velocity and tortuosity, which is not the case for head sweeps. (I,J,O) Quantification of the cool-mid (green) and mid-warm (orange) behavioural changes in relation to each species’ overall differences (purple). Black dots represent the mean, and the bars represent the first and third quartiles. The Oriental clade species have the largest overall changes in velocity and tortuosity across temperature zones. Except for *D. suzukii,* most of these changes occur within the cool-mid transitions. Changes for these same behaviours among the *Dmel*-subgroup species are more consistent across temperature transitions (closer or overlapping green/orange distributions). Other than *D. lutescens*’ and *D. takahashii*’s large reduction in head sweep size, clear clade differences for head sweeps were not observed.

Because our tracking of individual larvae allowed us to quantify speed, tortuosity, and head sweep sizes in relation to temperature gradients, we next asked if these elements of larval navigation reflect species’ preference differences that are consistent with our estimates based on temperature Electivity (above). We hypothesised that there would be significant behavioural differences between the species of the cooler-preferring Oriental group and the species of the *Dmel-*subgroup. In addition, our Electivity estimates led us to expect that, because *D. santomea* larvae are cool preferring, this species would display navigational behaviours more like the Oriental clade than any other species in the *Dmel*-subgroup.

We began by examining velocity and tortuosity across three temperature zones in our arena: cool (17.00-19.67°C), mid (19.67-22.33°C), and warm (22.33-25.00°C). As expected, the two measures are negatively correlated for all species (Pearson’s *R* ranged between -0.45 to - 0.84; Fig. S3), demonstrating that when larvae move faster their path is straighter^71^. Comparisons of speed and tortuosity between clades revealed that the Oriental species move faster and straighter across the three temperature zones compared to the *Dmel-*subgroup species (Wilcoxon rank sum test: *p*<0.001 for all temperature zones for velocity and tortuosity; Fig. 2D, E). On average, therefore, the Oriental species’ locomotion is faster than the species from the *Dmel*-subgroup clade. However, the magnitude of the differences between the two clades varied across temperatures, with the largest differences occurring within the warm and intermediate zones (Fig. 2D), suggesting species within the Oriental clade respond more aversively to the warmer temperatures than the *Dmel*-subgroup species. To investigate this further, we fit linear models to the data and asked if individual species within the Oriental group displayed stronger changes in behaviours in response to temperature transitions compared to the *Dmel*-subgroup species. Consistent with our expectations, we found that the slopes of the fitted regression models for both speed and tortuosity are significantly different between clades, with the species of the Oriental clade displaying significantly more rapid deceleration and increased tortuosity in response to cooler temperatures (see Table S3 for full stats; Fig. 2E,F). Grouping the species according to the trend of their responses separated the two clades and also highlighted *D. lutescens* and *D. santomea* - the species that we identified as having evolved the strongest cool preferences - as having the strongest responses within their respective clades (Fig. 2G,H).

The species variation in velocity and tortuosity prompted us to further examine how the magnitude of behavioural changes differed across the temperature zone transitions. We estimated the maximal velocity and tortuosity differences that were observed for each species between the cool and warm zones by taking the difference between randomly sampled velocity (or tortuosity) values between the two. To estimate the contribution to these maximal differences by the cool-mid and mid-cool transitions, we repeated the same sampling procedure between each of the two zones (Methods). Plotting these values accentuated the differences between the *Dmel*-subgroup and Oriental clade. Species from the Oriental Clade had the largest speed and tortuosity differences between the cool and warm zones (Fig. 2I,J), and, with the exception of *D. suzukii,* the behavioural differences evoked between the mid-cool temperature provided the bigger contribution. This pattern differed for the *Dmel*-subgroup species, for which the behaviours changed relatively consistently across the two temperature zone transitions (Fig. 2I,J). Together, these results provided additional evidence that the cooler temperatures elicit stronger attractive behaviours among the Oriental clade species compared to the *Dmel*-subgroup species and identified the responses to the mid-cool transition as the primary source of the differences (19.67-22.33°C to 17.00-19.67°C).

Analogous analyses of head sweeps in response to temperature zone transitions uncovered fewer differences between the Oriental and *Dmel*-subgroup species than we observed for velocity and tortuosity, consistent with previous findings^40^ (Fig. 2K-O). And though a linear model resulted in a significant negative relationship between the two (Fig. 2M, N; Stats in Table S3), it explained very little of the variation (adjusted *R*^2^ = 0.0159). The size of head sweeps is, therefore, significantly more variable over non-noxious temperature gradients compared to velocity and tortuosity. The overall variation in head sweep metrics between and within species was large. Despite this, investigation of the magnitude of species’ differences across temperature zones did reveal *D. lutescens* and *D. takahashii* to have the largest maximal reduction in head sweep size (between the warm and cool zones), consistent with their relatively strong cool preference (Fig. 2O).

### Agent-based modelling of species thermotactic differences

We have identified between-species differences in thermotactic behaviours based on both broad and fine-scale metrics. These changes raise questions about evolved differences in the larvae’s nervous systems. For *D. melanogaster*, considerable advances have been made in understanding the neural circuitry underpinning its homeostatic temperature preference, and so we next sought to leverage these insights together with an agent-based simulation approach to further examine species differences.

*D. melanogaster* larvae detect changes in innocuous cool and warm temperatures with two distinct peripheral neuron populations - Cooling Cells (CCs) and Warming Cells (WCs) - that express partially overlapping ionotropic receptors. CCs express Ir25a, Ir93a and Ir21a^72,73^ while WCs express Ir25a, Ir93a, and Ir68a^40^. Both neuron populations mediate avoidance behaviour to temperature changes, CCs specify avoidance to cooling and WCs specify avoidance to warming (Fig. 3A). Using behavioural, connectomic, and manipulative experiments, Hernandez-Nunez et al.^40^ also identified cross-inhibition between CCs and WCs, such that the activity of the cooling circuit inhibits the activity of the warming circuit and *vice versa*. In *D. melanogaster* larvae, it was found that cooling avoidance is initialised below 24°C and warming avoidance above 24°C. At temperatures close to 24°C, the two populations suppress avoidance behaviours, thereby establishing *D. melanogaster*’s homeostatic temperature preference at 24°C^40^.

**Figure 3:**
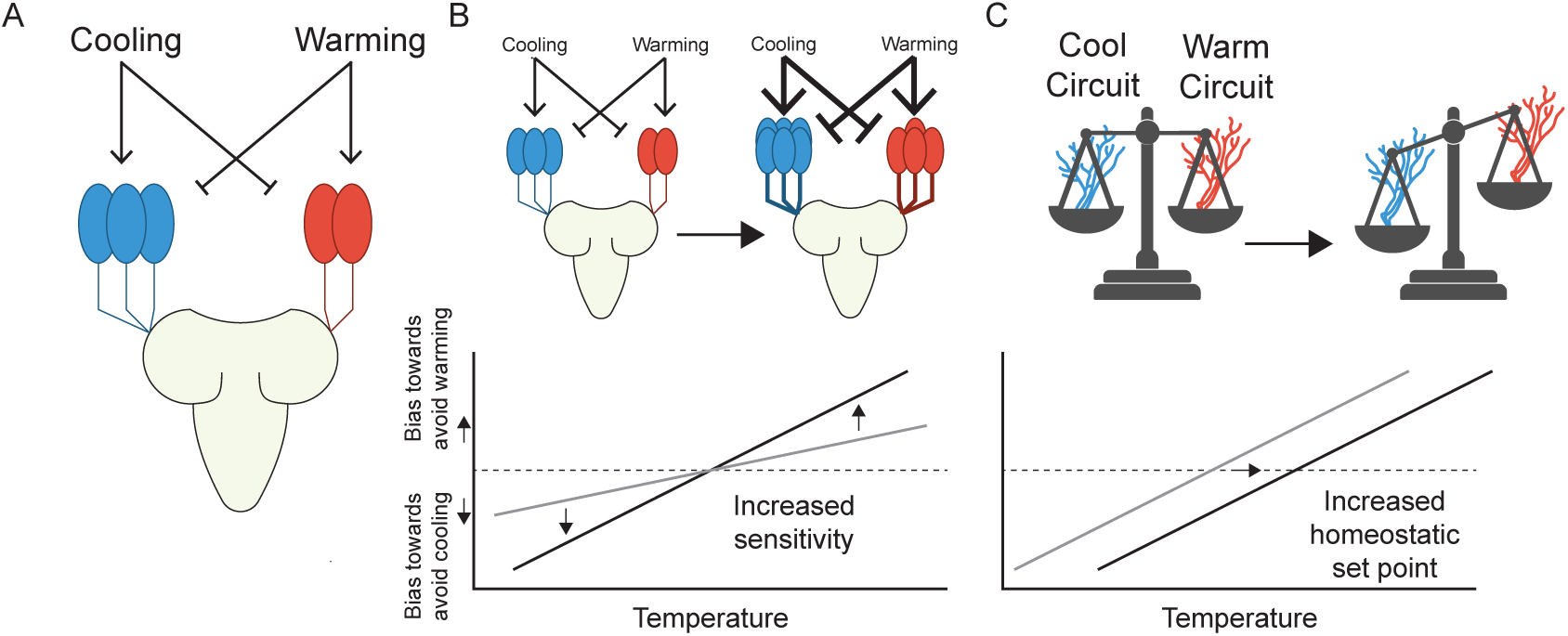
Diagram of larval cooling and warming circuits in the context of our simulation parameters. (A) Simplified representation of the three CCs and two WCs found in one side of the larval dorsal organ. Warming activates WCs and inhibits CCs and cooling activates CCs and inhibits WCs. Cross-inhibition occurs between CCs and WCs. Signals from these cells are transmitted to higher brain centres, ultimately influencing the behaviour that leads to temperature preference. (B) The slope parameter in the model effects the sensitivity of larvae to temperature. Larger slope values result in stronger avoidance behaviours at smaller changes in temperature. This could be driven by changes in the periphery such as changes in cell number (as depicted in the cartoon) or due to functional differences in the thermosensors expressed in cells, causing larvae to be more sensitive to changes in temperature. In the graph this is represented by the slope of the line, with larger slopes indicating higher sensitivity. The dashed horizontal line indicates where the cool and warm avoidance circuits are balanced. The further the line is from the dashed line, the more likely a larvae will turn back towards the homeostatic set point. (C) The set point occurs where there is an equal balance between the cool-avoidance and warm-avoidance circuits. On the graph, this is at the point where the larval avoidance response line crosses the dashed horizontal line. Shifts in the balance between the cool-avoidance and warm-avoidance changes the temperature of a larva’s homeostatic set point.

In Hernandez-Nunez et al.’s cross-inhibition model, the CCs and WCs dynamically generate neural responses upon detecting temperature changes. The relationship between the neural activity and avoidance behaviour (turning rate) is modelled using an empirically informed filter (See Equations 1 to 4 in ^40^). To introduce variability in the amplitude of neuronal responses at different temperatures, weights on CCs and WCs were introduced as free variables. These weights are temperature-dependent and modulate the magnitude of the neurons’ influence on turning rate and the strength of cross-inhibition on the other neuron type. For example, at temperatures below the homeostatic set point, the CCs are weighted to have a larger influence on turning rate. This results in larvae that are more likely to turn when going down the gradient and, due to cross-inhibition, less likely to turn when going up the gradient. Similarly, WCs are weighted to have a stronger influence on turning rate when going up the gradient when above the homeostatic set point. We reasoned that our large dataset could be used to fit species-specific parameters to this model, thereby providing a complementary approach for understanding how species differences may arise, and for generating hypotheses about their causes^64,74,75^.

Following the details established in the above cross-inhibition model we configured thermosensing virtual larvae and tested them using agent-based simulations^64^ in a simulated thermal arena that matched our experimental gradient. We implemented this model in Larvaworld, a recently developed behavioural analysis and modelling platform that supports a broad range of agent-based larvae simulations, extending it to integrate thermotactic behaviour^75^ (Methods). To estimate the species-specific aversion parameters (lateral body-bending behaviour in response to temperature changes), we simulated larvae over a grid of “homeostatic set points” and “slope values”. The homeostatic set points are the temperatures at which the weights for the WCs and CCs are equal and the slope values relate the weights to temperature (see Equations 2 - 5 in Methods; Fig. 3B, C). A higher slope elicits a stronger aversive behavioural response in simulated larvae as they move away from the homeostatic set point. We made the simplifying assumption that the relation between temperature and behavioural bias for the cooling and warming cells are linear and symmetric with a scalar slope parameter, which significantly reduces the dimensionality of our simulations (Methods). Estimates of these two model parameters were acquired by applying rejection sampling to the simulated datasets (Fig 2A-H; Table S4; simulated agents per grid point = 1000; acceptance threshold = Euclidean distance < 1.75 compared to our empirical data; Methods).

Examination of the posterior distributions of the model’s parameters revealed variation in the best-fitting point estimates for slope and set point across the eight species (Fig. 4A-H white circles, see Table S4 for best-fit parameter values). The estimated set points are consistent with our empirical measurements of Electivity (above) and fall within the individual empirical *E*_breadth_ boundaries for all species, with a tendency to lie close to the higher end of the empirical ranges. Inspection of the 95% credible interval of the joint parameters (Fig. 4I, J), revealed significant differences in the set point among multiple species while slope values largely overlapped. These results suggest that shifts in the balance of signals from the cool- and warm-detecting circuits drive species differences rather than changes in sensitivity to cooling or warming temperatures. Our inspection of a sparser grid of simulations over a larger range of slope values revealed that lower slope values were better for predicting our empirical data for all species, indicating that larval exploration in this temperature range is driven by relatively weak aversive behaviours (Fig. S5). This is perhaps not surprising as the temperature range is innocuous and any thermal preference may be minimised by foraging needs or other behavioural drivers. Despite the weak aversive behaviours, it is notable that species differences in the homeostatic set point values – themselves indicators of temperature preference – could still be clearly identified.

**Figure 4:**
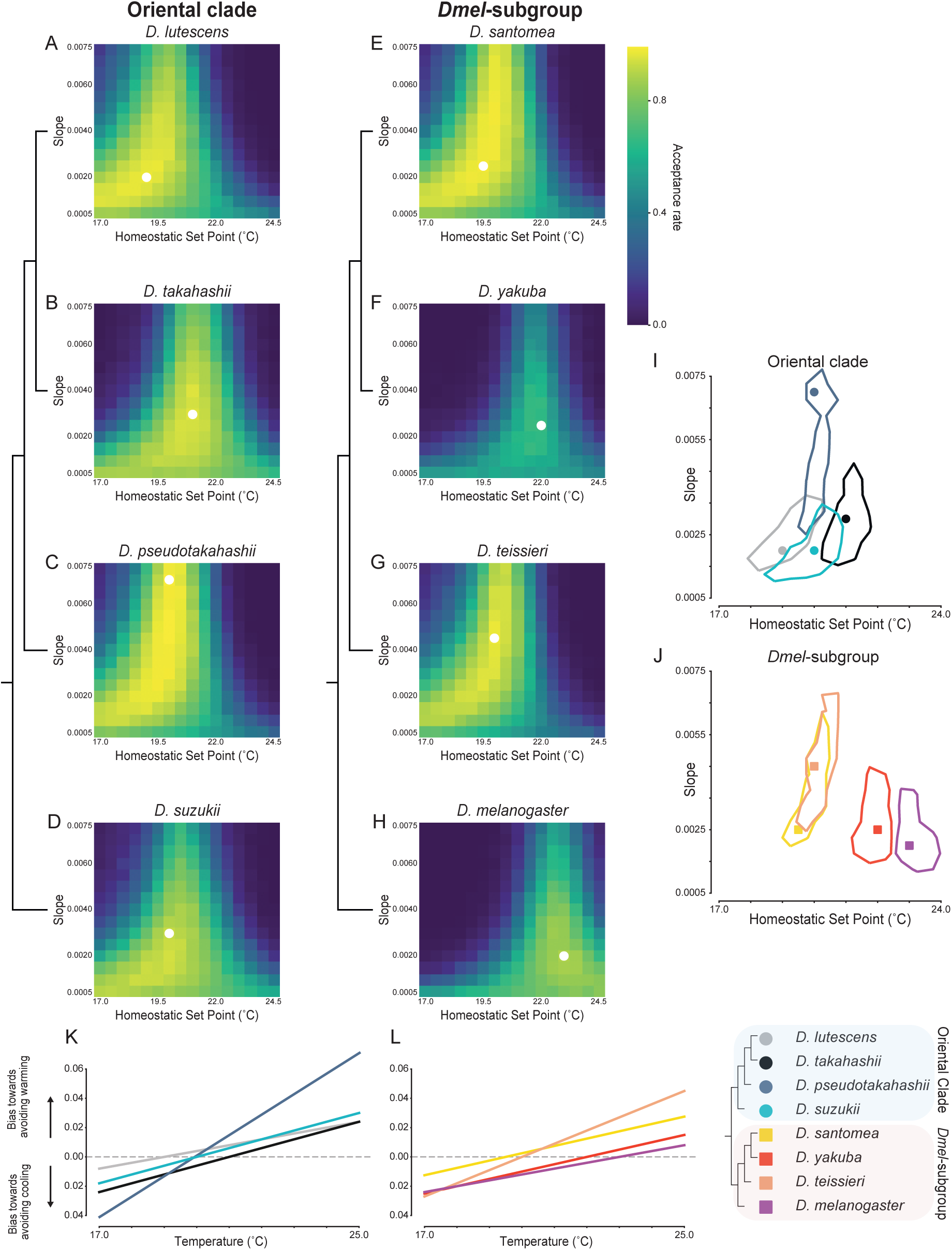
Agent-based simulation results indicate that shifts in the balance of cool and warm circuits, not sensitivity, drives species specific temperature preference profiles. (A-H) Heatmaps displaying the acceptance rates of simulations for each parameter combination, each containing a set of 1000 simulated agents. The simulations were run with two variable parameters, homeostatic set point (x-axis) and slope values which represent sensitivity of circuits (y-axis). If the best fitting simulations have a low temperature and high slope, this indicates that the species has a high affinity for cooler temperatures. The white dot represents the best fitting simulation for the species. (I-J) Contour plots displaying the parameter combinations that are within the 95^th^ percentile of best fitting parameter sets, with the best fitting simulation denoted with a circle/square. In (I) the Asian clade species all tend to have best fitting simulations towards the cooler end for their homeostatic set points, with *D. lutescens* and *D. takahashii* still having a clear difference. *Drosophila pseudotakahashii* and *D. suzukii* are intermediate between the two. The *Dmel*-clade (J) have a larger spread across the homeostatic set point temperature. *Drosophila santomea* and *D. teissieri* show similarity in their best fitting simulations, whereas *D. yakuba* and *D. melanogaster* match simulations with a higher temperature preference, representative of the temperature preference difference between the two pairs of species. (K-L) The bias of larvae moving away from cooling (below zero) and warming (above zero) according to best fitting model parameters. The further from zero the line is, the stronger the contribution from the warm cells (above zero) or the cool cells (below zero). The point where a line crosses zero is the homeostatic set point (warm and cool circuits have balanced contributions to the avoidance behavioural output). For reference, the species topology is shown to the right.

Within the Oriental group, we found the largest difference (2°C) in set point was between *D. takahashii* (21°C, Fig. 4B) and *D. lutescens* (19°C, Fig. 4A). The lack of a significant overlap in the joint distribution of the model parameters for this pair of species highlights their divergence in temperature preference. Inspection of the parameters that best fit *D. takahashii*’s avoidance below 20°C revealed stronger avoidance to cooling at these temperatures compared to *D. lutescens* (Fig. 4K). At temperatures above 21°C, both species exhibit a similar avoidance behaviour (with equal avoidance to moving up the gradient at 25°C; Fig. 4K), consistent with the observation that neither species explored temperatures above 22°C in our behavioural assays. *D. suzukii’s* and *D. pseudotakahashii’s* estimated homeostatic set points, 20°C for both, were intermediate to *D. lutescens’ and D. takahashii*’s. Although the joint distribution of their model parameters shows significant overlap (Fig. 4C, D), *D. pseudotakahashii*’s best-fitting slope value is considerably higher than *D. suzukii*’s – consistent with *D. suzukii’s* shallower *E*_peak_ profile when compared to *D. pseudotakahashii*’s (Table 1).

In the *Dmel*-subgroup, *D. santomea* and *D. teissieri* share similar posterior distributions (Fig. 4J), but differ significantly from *D. yakuba* and *D. melanogaster*, which in turn have significantly different distributions from each other (Fig. 4J). *Drosophila melanogaster* was found to have the warmest overall predicted homeostatic set point at 23°C (a close match to Hernandez-Nunez et al.’s 24°C) and is the only species whose posterior distribution does not overlap with another’s. When inspecting the avoidance behaviour weights from the best-fitting simulations (Figure 4L), we observed that *D. melanogaster* and *D. yakuba* avoid temperatures below 20°C with similar avoidance profiles. At temperatures above 22°C, however, *D. yakuba* is more likely to turn down the gradient than *D. melanogaster*. The two cooler-preferring species in the clade, *D. santomea* and *D. teissieri*, were found to have a significantly stronger tendency to avoid the warmest temperatures of our assay (Fig. 4L).

Overall, our agent-based simulation approach demonstrates that the simple cross-inhibition model can be used to parameterise species-specific differences in temperature behaviour. Despite using larva tracks collected over non-noxious temperatures, we were still able to infer significantly different model estimates among members of the Oriental clade and the *Dmel*-subgroup clade, including *D. lutescens*–*D. takahashii* and *D. santomea*–*D. yakuba* - the species pairs that have evolved cooler preference in parallel. Importantly, the differences that we found suggest that changes in sensitivity to either warm or cool temperatures have remained unchanged and instead evolutionary shifts in the balance of signals from the cool- and warm-detecting circuits drive species temperature preference differences.

## Discussion

Understanding how animals adapt (or fail to adapt) to novel environments is a fundamental biological question, and in the context of novel thermal environments also presents pressing challenges due to rapid climatic changes^76,77^. To understand how temperature-related behaviours evolve in poikilotherms, animals that are particularly vulnerable due to their exclusive reliance on behavioural strategies for thermoregulation, we have carried out a large-scale comparative thermotaxis study using *Drosophila* larvae from diverse thermoecologies. By applying phylogenetically informed analyses to thousands of tracked individual larvae across eight closely related species, we were able to identify recurrent changes in temperature preferences that have evolved over short and intermediate timescales.

Evolution in temperature preference based on aggregated larval locomotory tracks revealed that temperature preference evolved recurrently within and between the Oriental clade and *D-mel* subgroup, two closely related clades from the *D. melanogaster* species group. Species from the Oriental clade tend to prefer cooler temperatures (Fig. 1D), and we inferred a “cool shift” in their common ancestor. However, given the fast rate that temperature preferences evolve (Fig. 1D), additional outgroup species are needed to better inform this transition. Temperature preference difference based on individual larva’s velocity and tendency to turn (tortuosity) along thermal gradients also support the rapid evolution of thermotaxis. Species from the Oriental clade avoid the warmer temperature zones (19.67-25°C) by exiting them more quickly and with fewer turns than species from the *Dmel*-subgroup. Notably, the Oriental clade has evolved to be faster crawlers irrespective of temperature (Fig. 2D). While velocity is related to other phenotypes, we hypothesise that this is an adaptation resulting in part from their narrower thermal preferences and elevated demands for rapid thermotactic responses.

Further underscoring the rapid evolution of temperature preference is the discovery of a parallel divergence between the larvae of the two most closely related species-pairs, *D. takahashii*-*D. lutescens* and *D. yakuba*-*D. santomea* (Fig. 1D). In both instances, there was a behavioural match to the species’ known thermoecology, with the species that live in the cooler habitats (*D. lutescens* and *D. santomea*) found to prefer a cooler temperature range compared to their sister species that live in warmer habitats (*D. takahashii* and *D. yakuba*; Fig. 1). Measurements of the individual larva’s velocity and tortuosity provided additional support for the recently evolved differences, with *D. lutescens* displaying the largest avoidance responses to warm temperatures within the clade (Fig. 2M,N). Though temperature-related behaviours have not been previously studied within the Oriental clade, thermotolerance experiments were motivated by the cooler northern distribution of *D. lutescens* in comparison to the warmer subtropical distribution of *D. takahashii*. These studies found *D. lutescens* to be more cold tolerant compared to *D. takahashii* across all life stages^30,54^. For example, the temperature leading to 50% experimental population mortality of adult flies is 4.5-4.6°C for *D. takahashii* while it is 0.5-1.1°C for *D. lutescens*^31^. Although thermotolerance differences may not be indicative of temperature preferences within an innocuous range, this question remained open. Our results have demonstrated that *D. lutescens* indeed prefers significantly cooler temperatures than *D. takahashii* (*E*_peak_: 17.0°C compared to *E*_peak_: 19.7°C; See Fig. 3C and Table 1). Together, these observations are consistent with the thermotactic changes evolving adaptively and motivate future fitness assays carried out for the two species over a similar range of temperatures.

Within the *Dmel*-clade, *D. santomea* adults have a behavioural preference just below 23°C whereas *D. yakuba* adults have a higher temperature preference of 26-27°C^51,52^. We have shown that a difference in temperature preference of ∼2-3°C also exists between these species at the larval stage (*D. santomea E*_peak_ = 18.5°C, *D. yakuba E*_peak_ = 20.3°C). Electivity-based differences were again supported by individual larva’s velocity and tortuosity, with *D. santomea* displaying the strongest avoidance to warm temperatures within the *Dmel*-subgroup (Fig. 2). As with *D. lutescens*-*D. takahashii*, we argue that our observations for *D. santomea* and *D. yakuba*, in combination with previous fitness and tolerance assays, are consistent with recent adaptation to different thermal environments. The consistency in temperature preference between adults and larvae also extend to *D. teissieri*, the closest outgroup species of *D. yakuba* and *D. santomea.* Adult *D. teissieri* have temperature preferences comparable to *D. santomea*, ∼23°C^51^. We likewise found that the temperature preference of *D. teissieri* larvae is more similar to *D. santomea* (*E*_breadth_ = 17.0-20.6°C compared to *E*_breadth_ = 17.0-20.5°C) than to *D. yakuba* (*E*_breadth_ = 19.4-22.1°C) (Fig 1D and Table 1). These results are therefore consistent with our understanding of the species’ thermoclimatic ranges: *D. santomea* is found at cooler higher altitudes than *D. yakuba* on the island of São Tomé^48,52^, and even though there is considerable range overlap between *D. yakuba* and *D. teissieri*, the latter tends to occupy cooler mid-high elevations within this range^51^.

As most, if not all, of the eight species that we studied contain considerable genetic diversity^78–81^ it is expected that they display behavioural polymorphism too. To guard against mischaracterising a species’ thermal preference due to a single outlier strain, we analysed three strains per species (except *D. pseudotakahashii* for which we were able to obtain only a single strain). We found significantly more variation in thermotaxis between species than within (ANOVA on *E_peak_* : *F*-value = 6.01, p-value <0.01), indicating a significant portion of this behaviour is heritable and that we have quantified genetic divergence and not plasticity. Interestingly, the most behaviourally variable species was *D. melanogaster*, which also has the warmest temperature preference (*E*_peak_ = 21.4°C). However, inspection of the full Electivity profile reveals that the preference is weaker than in other species (*E*_peak_ median: 0.026), resulting from between-strain variation in Electivity (Fig. S2-3, Table S1). Previous work on *D. melanogaster* w1118 larvae reported a preferred temperature for third-instar larva of 24°C^82^. We also included w1118, and estimated a comparable *E*_peak_ of 22.8°C. However, the other two strains, Canton-S and a Chinese strain (B63^79^) had a notably different *E*_peak_ of 19.1 and 20.8°C, respectively. These differences suggest considerable thermotaxis variation across *D. melanogaster* populations which are genetically structured^79^ and serves as a reminder that phenotypes measured from w1118 - a common lab strain used in many behavioural assays - may not be representative of the species. We hypothesise that the lack of strong temperature preference in *D. melanogaster* may have contributed to this species’ ability to successfully inhabit the globe. An interesting contrast, however, is the agricultural pest, *D. suzukii*, which is currently undergoing a global expansion^60,83,84^ but has a relatively strong cool preference (*E*_peak_: 17.4°C; *E*_peak_ median: 0.149), consistent with previous reports in adults^85,86^. This *D. suzukii* example implies that a weak temperature preference does not necessarily precede a poikilotherm’s global spread.

How might thermotaxis evolve? Using simulations based on the neural circuitry that drives *D. melanogaster* larval thermotaxis behaviour^40^, we were able to explore how this circuitry could evolve between species. We investigated two non-mutually exclusive hypotheses for how thermotaxis might evovle: (1) a change in thermosensitivity feeding the avoidance pathways and (2) shifts in balance between the Cooling Cells (CC)/Warming Cells (WC) circuits. The former could entail changes in the peripheral cell’s (CC or WC) sensitivity, number, or morphology^87^ . The latter does not necessarily rely on sensitivity and could be attributed to changes in upstream circuitry that influences the balance between the two avoidance circuits, causing shifts in homeostatic set points. In our simulations, varying both the sensitivity and the balance between the circuits did not result in evidence for differences in sensitivity; we found that sensitivity of larvae towards the temperatures tested (18-25°C) does not vary greatly between species and that it tended to be relatively low (Fig. 4I,J, S4). This indicates that within the innocuous range of temperatures that we tested, none of these species differ in their strength of avoidance to perceived changes in temperature, nor do they differ in their preferences towards their homeostatic set points. Instead, we found evidence for the differences between species being largely driven by differences in the balance between the CC and WC circuits.

### Limitations of the study

#### Species sampling

An important result of this work has been to reveal how quickly larval temperature preferences evolve. An upshot of its rapid evolution and multiple species-specific changes is that it limits the ability to polarise when several of the preference shifts occurred. Additional species sampling would provide finer resolution. In particular, additional outgroup species would help to polarise the cool/warm preference that differentiates the Oriental clade and the *Dmel*-subgroup.

#### Behavioural Assays

The design of our thermal arena prevented temperatures below 17°C from being held stably. As a result, we have underestimated *D. lutescens’ E*_peak_ and its lower bound of *E*_breadth_. Our results for this species are therefore conservative.

#### Simulations

Previous models using *D. melanogaster* larvae have shown that turning rate variation alone is enough to predict taxis behaviours^64^. For this and other reasons, they assume that larvae move at the same speed across temperatures. However, we demonstrated that larval velocity does change across temperatures and that species have different speeds. We propose that adding model flexibility for velocity changes will provide better matches to empirical data and that future developments of the model will benefit by adding speed as a variable.

The cross-inhibitory model was previously parameterised using empirical data from *D. melanogaster*^40^. Analogous data does not exist for other species and so we made the simplifying assumptions that the weights are linear and symmetric between the WCs and CCs. Although the dimensionality of the simulations would quickly become prohibitive, varying the weights of the CCs and WCs individually and introducing more complex weight-temperature relations may likewise provide better matches to empirical data.

## Methods

### *Drosophila* species, maintenance, and larvae collection

All species used in this study belong to the *D. melanogaster* species group. Each species was studied using three strains (apart from *D. pseudotakahashii*, where only one strain was available). The species and strains that were used in this study are shown in Table S5. All species were maintained in vials containing a standard fly media composed of yeast, agar and cornmeal supplemented with Formula 4-24 Instant Drosophila Medium, Blue (Carolina). Flies were kept at 23°C in a 12:12 light cycle.

To collect third-instar larvae across the different species, we first tested the developmental times required to reach this stage at 23°C. Limiting a window of time for egg laying to two hours, we found that for *D. melanogaster*, *D. yakuba*, *D. takahashii*, *D. suzukii,* and *D. teissieri*, five days were needed, while for *D. lutescens*, *D. pseudotakahashii*, *D. santomea* the duration was six days. On the day of an experiment, larvae were floated in a 15% sucrose solution and third-instar were collected and rinsed with tap water. We recorded the approximate number of larvae applied to the arena prior to starting the assay and we determined the final sample sizes based on the larval tracks kept after filtering steps.

### de novo D. lutescens genome assembly

Seven of the eight species used in this study had reference genomes available (*D. santomea, D. yakuba*, *D. melanogaster*, *D. teissieri*, *D. takahashii*, *D. pseudotakahashii* and *D. suzukii*). We additionally generated a *de novo* assembly for *D. lutescens*. We collected 200 *D. lutescens* AK96-3 male flies and prepared them for DNA extraction by flashing freezing flies and rupturing cells with metal beads in a cryomill. We then used the Qiagen DNA extraction kit to extract long DNA strands, followed by gentle shaking in a cold room (4°C) for two weeks to dissolve DNA in a buffer. Library preparation and sequencing on two lanes of PacBio’s SMRT cell V2 was done by the Lausanne Genomics Facility.

The raw PacBio reads were assembled and subsequently used for a single iteration of polishing using Flye^88^. Heterozygous contigs were assigned as haplotigs, and contigs with extremely low or high coverage were assigned as artefacts using PurgeHaplotigs^89^. The genome was polished using RNAseq reads with two rounds of Pilon. RNAseq reads from *D. lutescens* whole bodies were generated using the same methodology as Bontonou et al.^90^. Alignment required for this polishing was done with STAR’s 2-pass mode^91^. The Sequence data used for the *D. lutescens* assembly is available on GenBank under BioProject PRJNA1002970.

### Species divergence estimation

To obtain single copy orthologues to build a phylogeny we used OrthoFinder^92^. This required all genomes to be soft-masked. We built a *de novo* repeat library per species (+ 12 other genomes to aid in calibrating node dates downstream) using RepeatModeler2.0 with the LTRStruct flag^93^. The library was combined with Dfam3.0 as a custom species-specific database on RepeatMasker^94^, to soft-mask the genome. We then annotated the genome using the BRAKER^95,96^ pipeline with evidence from the Arthropoda orthologue database (v10)^97^, and for *D. lutescens* we also included the RNAseq data. Orthofinder was then ran with the following flags: -M msa -T fasttree. The resulting species tree from OrthoFinder was then input into MEGA11^98^ to date the tree using secondary calibrations based on node estimation dates from^99^ These are shown in Table S5. The resulting tree (Fig. S5) was then pruned using ape in R.

### Arena construction

The arena was built using a 170×170×0.5mm aluminium plate placed on top of four Peltier elements (Fig. 1B). The temperature of the Peltier elements was controlled by an Arduino microcontroller. We employed a closed feedback loop to achieve our desired temperature range, where temperature sensors placed on the aluminium plate directly above the Peltier elements, provided real-time temperature data to the microcontroller. This could then modify the power provided by the bench top power supplies (PeakTech) to the Peltier elements, using pulse width modulation, until the desired temperatures were reached. To prevent larvae from escaping, the arena’s perimeter was surrounded by a thermal wall, which contained a nichrome wire maintained at a noxious temperature range of 50-60°C (the wire was not in contact with the gel surface and inspection of our thermal imaging indicated the wire had no discernible impact on the temperature of the arena’s surface).

The build also consists of two cameras that record the arena from above. A camera to record larvae exploring the arena, and a FLIR thermal camera to record the thermal gradient. Illumination for the camera was provided by red LED lights, which should not influence larval behaviour as larvae lack photoreceptors to light in the red range^100,101^. To prevent external disturbance from light, wind, and sound, we encased the arena with an outer shell made of cardboard and black fabric. Details of the arena build are available on https://gitlab.com/EvoNeuro/patchythermalgradient.

### Running behavioural assays

All assays were carried out on a 170×170×6mm 4% agarose gel, which was placed upon the aluminium plate. To provide contrast 1% charcoal was added, along with 10% sucrose to encourage larvae to stay on the arena. To reach our desired temperature range of 17-25°C, we set the temperatures of the arena to be 15°C on the cold sides, and 29°C on the hot sides, accounting for the difference in temperature from the Peltier elements to the top of the assay gel. To run an assay, floated and rinsed third-instar larvae were brushed onto the middle of the arena, and were allowed to explore for twenty minutes, whilst being recorded. Assays were conducted in a dark room with 19°C ambient temperature between 15 June to 04 August 2021. To limit external biases, the arena was rotated by 90° every two weeks, changing the positioning of the cold and hot temperature sources.

### Image processing

We used the cameras propriety software Spinnaker SDK to save TIFF images at 10Hz onto a Dell Precision 3640 computer. The thermal camera data was saved using a Python script that read data from the thermal camera, a modified version of uvc-radiometry.py (https://github.com/RDelg/Footshot/blob/master/uvc-radiometry.py), set up to capture the thermal gradient topology three times per minute. Due to problems with the thermal camera’s internal heating, we had to smooth abnormal spikes in recordings using an in-house Python programme (see script smoothspikes.py).

A quality control step was run with the following criteria: 5% of the arena had to be below 17.5°C and above 24.5°C, and this had to be maintained during over 90% (91.67%) of the run. Additionally, no pixel on the arena could fluctuate more than 3°C (see script QC_check.py). Image data from the camera and thermal camera, from assays that passed the quality control steps, were cropped to contain only the arena using a custom Python script with opencv2 (see script click2crop.py).

### Track analyses

Cropped camera data was input into the larval tracking software FIMtrack^102^ to obtain coordinates for larval movement during the run. To reduce file size, we ran an awk command to remove tracks shorter than ten seconds (see script 02cleandata.sh). Tracks were classified as non-moving if they did not travel more than 0.5mm accumulatively and 0.3mm from their origin (see script 03showtracks.py) and were subsequently manually removed with our deletetracks.py script. Clashes between larvae during runs caused loss of larval identity, splitting their runs into multiple tracks. We automatically joined tracks from clashes using an algorithm that detects when two tracks abruptly end on the same frame, and joins them to the reciprocally closest track, in terms of time and distance (with time taking priority; see 04jointracks.py with -ac flag to automatically join clashes).

The remaining disjointed tracks lost due to problems in tracking were resolved using a similar joining algorithm. Tracks were joined if they were reciprocally the track that ended and started the closest in time and distance. This was run in multiple rounds, with the first round requiring the end point of the first track and the start point of the second track to be within 150px and 22.5s of each other. This was run iteratively until no more tracks could be joined. Subsequent rounds became less stringent, with the second round distance being extended to 500px and end to start timing being increased to 75s of each other, and in the final round all restrictions were dropped. In rounds one and two, we also placed a restriction on joining if the speed of the larva to reach end point of the first track and the start point of the second track was deemed unreasonable (round 1 <15px/s and round 2 <30px/s).

Complete larval tracks were then matched to temperatures using the thermal images that were taken closest in time to that point. We removed the first two minutes of every assay as a burn in period, allowing larvae to acclimatise to the assay.

We used a modified version of Ivlev’s Electivity (*E*) as our temperature preference index (Equation 1)^58^. We calculated this for 1°C windows with 0.1°C steps and described a larva’s temperature preference profile by the temperature of the maximum *E* value (*E_peak_*) and the range of temperatures where they spent time (*E_breadth_)*.

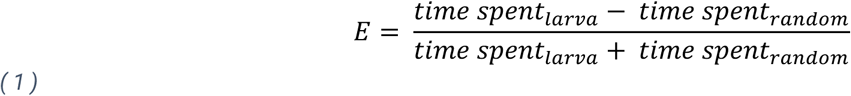

To create null tracks (*time spent_random_*) a set of 1000 randomly moving agents was simulated for each run on the same temperature gradient (see script 05nulldistribution.py). The average speed and turn rate of the larvae of that run were input as parameters for the simulated agents, which were calculated by the 04autojoin_tracks.py script with the -ndp option. Starting positions were randomly generated within the middle 33% of the arena, and starting orientation was also random. Simulated agents explored the arena for twenty minutes, and the first two minutes were removed for the simulations as we did for actual runs.

### Agent-based simulations

The description of the cross-inhibition model for larval thermotaxis in *D. melanogaster* prompted exploration into how parameters of the model differ for other species. Within the model weights describing the influence of Cooling and Warming Cells to causing avoidance behaviours with respect to temperature were estimated for *D. melanogaster* empirically. We aimed to estimated how and if these weight parameters differ between species.

For our simulations, we used the software package <u>Larvaworld</u>^75^. Larvaworld supports several sensory modalities such as olfaction, touch and wind mechanoception. For each modality the respective sensors are available when configuring a virtual larva’s behavioural architecture and the respective sensory landscape is available to superimpose onto the virtual arena, eventually allowing for closed-loop sensorimotor simulations. We therefore extended the platform by implementing thermosensation, enabling thermal gradients across the arena (thermoscape) and introducing thermosensing agents. Gradients are created by setting a baseline plate temperature and placing cold/hot sources on the plate that modify the base temperature through a Gaussian spread. In our case, we set this to a 17×17cm gradient, with the four temperature sources located at the same position as in the original experiments (plate temperature at 21°C, two cold at 14°C, two warm at 28°C, Gaussian spread: 0.1 with SciPy’s multivariate_normal function).

Virtual agents have thermosensors located at the tip of their heads by which they dynamically detect temperature changes. While each sensor can vary in its thermosensitivity, they all converge to form a single locomotion-influencing input that biases the larva’s turning behaviour towards positive or negative thermotaxis. We set the cool sensor’s gain so that its activation encourages turning when moving towards cooler temperatures and inhibits turning when going up temperatures, whereas the warm sensor does the opposite, encouraging turning when going towards warmer temperatures and inhibiting turning when going down temperatures. Temperature-dependent modulation of turning is based on an earlier model, proposed in the context of chemotaxis, by Wystrach et al. (2016)^64^, as later extended and used in Larvaworld^103^.

Both sensors are always active, but their aversive strength (determined by gain in Larvaworld) is linearly weighted with absolute temperature (Equations 2 and 3). For example, the warm sensor encourages turning more strongly when going up the temperature gradient at warmer temperatures than at cooler temperatures and inhibits turning more strongly when going down the gradient at warmer temperatures than cooler temperatures. The cool sensor, on the other hand, has stronger aversive properties when going down the gradient at cooler temperatures than warmer temperatures.

In our simulations, the weight of each sensor was determined by the slope parameter of a linear function, with the cool sensor having a negative slope and the warm sensor having a positive slope. To reduce the number of parameters, both sensors were assigned equal slope values of opposite sign and limited values between 0 and 1. The weights of both sensors always overlapped at 0.5 and as the aversive properties of both sensors are very close around the corresponding temperature, it results in random movement at this temperature. Ultimately, this leads to a preference for that temperature, which can be referred to as the homeostatic set point. The weights of the cool circuit (*W_cool_*) and warm circuit (*W_warm_*) are calculated by the following formulae:

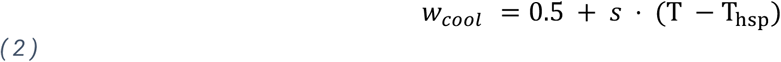

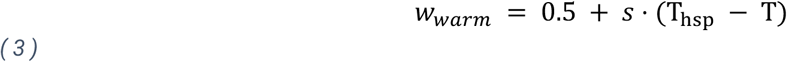

where *s* is the slope value (that determines sensitivity), T is the temperature where the agent is at, and T_,-._ is the temperature of the homeostatic set point. The two weights (*W_cool_* and (*W_warm_*) are equal when T = T_,-._.

To model the behavior-modulating signal (A_O_) that a larva extracts from its thermosensory environment, we assume that changes in thermal perception are proportional to the logarithm of changes in sensed temperature as dictated by the Weber–Fechner law^104^ widely used across sensory modalities. We add a decay term which gradually returns A_O_ back to zero. The equation is:

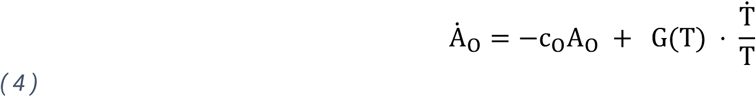

where c_O_ = 1 is a decay coefficient and G(T) a temperature-dependent gain parameter that is proportional to slope and the relationship between the homeostatic set point and temperature. G(T) is determined by the subtraction of the weights presented in Equations 2 and 3. The gain value (always set to below 0) determines the avoidance behaviour in Larvaworld.

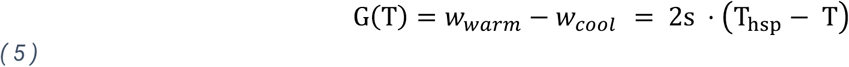

Overall, the larvae’s movement depends upon the interplay between the homeostatic set point temperature (T_hs*p*_) and the temperature the larvae is at (T), alongside the change in temperature over between steps (Ṫ). The parameter G signifies a set gain value that is what determines the avoidance behaviour in Larvaworld.

We varied the point of this overlap, the homeostatic set point, in our simulations. The first set of simulations was run with homeostatic set points ranging from 17-25°C (0.5° step size), and slopes were varied from 0.0025-0.05 (0.0025 step size). After finding that lower slopes were better fits for all species, we ran another set of simulations with the same homeostatic set point range, but slopes ranging from 0.0005-0.007 (0.0005 step size). Each simulation in the first round consisted of 500 simulated agents, and in the second round, this was doubled to 1000 virtual larvae. A set of 2000 “temperature-blind” virtual larvae (i.e. equal avoidance output of warm and cool circuits across all temperatures) were also simulated to calculate our temperature preference index (Equation 1).

### Determining temperature preference

Once Electivity was calculated across bins for every individual larva, we determined *E*_peak_ by the temperature with the highest mean Electivity. *E*_breadth_, the range of temperatures where larvae were comfortable, was determined using the sign test. The null hypothesis of this statistical test is that the median Electivity of a temperature bin is equal to zero. Temperature bins which resulted in statistical significance after multiple correction were considered strongly preferred temperatures if positive, and if negative they were considered aversive temperatures. Temperature bins with medians that were at 0 (not significant in the sign test) were labelled as preferred temperature zones, with the lowest temperature forming the lower boundary of *E_breadth_* and the highest temperature being the upper boundary of *E_breadth_*. As *E_peak_* is skewed towards negative values (due to multiple bins and constantly moving larvae, Electivities reach -1 frequently and it is rare to have Electivities closer to 1; Figure S1).

When comparing key species pairs, we used a Mann-Whitney U test due to the non-normal distribution of the Electivity values. The null hypothesis was that species were not different in their Electivities, and the alternative was testing if one of the species had a stronger preference. As we tested across temperature bins, these were corrected for multiple testing. To check if variation within species (between strains) was lower than between species, we ran a phylANOVA on *E*_peak_, using the phytools package in R. A significant ANOVA value indicates that there is less variation within species than between species, supporting the grouping of species. To check for shifts in *E*_peak_, and the upper bound of *E*_breadth_, we used the phylolm package in R and input our dated phylogeny. The lower bound of *E*_breadth_ was not analysed as there was little variation due to the lower limit of the temperature gradient (17°C).

### Analysing fine-scale behavioural metrics

We calculated velocity using the compute_velocity_window function in the custom script polarplotsV2.py, with window size set to 3s and each step being a frame (step size: 1/10^th^ of a second). We recorded velocity in both mm/s and body lengths/s. Tortuosity was also calculated using a window-based method over various window sizes (2,5,10,20,30 seconds). This is calculated by dividing the as-the-crow-fly distance moved by the larva in that time by the actual accumulated distance the larva travelled, then subtracting the resulting value from one. The average of all the windows is taken to give the final tortuosity value.

To detect head sweeps, we first pulled the body bending feature for every track provided by FIMtrack. We then used SciPy’s findpeaks function to detect head sweeps. Body bends greater than 30° that return to within 10° of a straight posture were considered head sweeps, with a buffer of 2 seconds between head sweeps. In the final analysis, only head sweeps greater than 45° were kept.

We measured correlation between speed and tortuosity using Pearson correlation coefficient, for each species. We analysed clade differences in speed, tortuosity and head sweeps using a Wilcoxon rank sum test. At the species level, we carried out linear regressions across temperatures. The function emtrends from the emmeans package in R was then used to compare trends between species, a compact-letter display was generated to group species after pairwise comparisons.

To measure differences in these navigational metrics at different temperatures, we split the arena into three temperature zones (cool: 17.00-19.67°C, mid: 19.67-22.33°C, and warm: 22.33-25.00°C). We took measurements of velocity, tortuosity, or head sweeps for each species in each zone, and subtracted this in a pairwise manner warm-cool, mid-cool and warm-mid, we did this 1000 times with random selections of the behaviour. We then performed a t-test to compare differences between different zones within species. We were able to also test for magnitude difference between species by using a t-test. All these analyses are available in 07_finerscalebehaviours_other.py.

### Analysis of agent-based simulations

We applied a rejection sampling approach to fit species-specific models to our data. For every agent in each simulation, we calculated Electivity using the “temperature-blind” larvae as our null set. To determine which of these simulations best fit each species, we compared them to the species’ mean Electivity using a distance measure (Euclidean distance) and rejected individual agents that exhibited distances larger than 1.75. For each species we then determined the 95^th^ percent credible interval (i.e. the 95th percentile of the 2D distribution), which we visualised using a contour plot (modifying the contour so that the whole of the best fitting simulation square was included, Fig. 4 I,J). The best fitting simulation was that with the highest acceptance rate (white circles in Fig. 4 A-H). To calculate how similar species were overall on simulations, we ran a PCA using the acceptance rates for all simulations per species (Fig. S6).

## Data, code and resource availability

All original code has been made publicly available on our lab’s repository: https://gitlab.com/EvoNeuro/patchythermalgradient. DOIs are listed in the Resources Table.

## Resource availability

### Materials availability

All fly strains used in this study are available from the lead contact upon request.

### Lead contact

Further information and requests for resources should be directed to and will be fulfilled by the lead contact, Roman Arguello

## Resources table

| REAGENT or RESOURCE | SOURCE | IDENTIFIER |
| --- | --- | --- |
| **Strains** |  |  |
| *D. lutescens* AK96-3 | Kyorin | E-12001 |
| *D. lutescens* NGN22 | Kyorin | E-12020 |
| *D. lutescens* 29 |  |  |
| *D. takahashii* KMM9 | Kyorin | E-12231 |
| *D. takahashii* IHYT1 | Kyorin | E-12230 |
| *D. takahashii* 39 |  |  |
| *D. pseudotakahashii* 0301.01 | Kyorin | E-24401 |
| *D. suzukii* WT3 | US Stock centre;<br>Benjamin Prud'homme |  |
| *D. suzukii* NGM-2 | Kyorin | k-awa036 |
| *D. suzukii* | Bruno Lemaitre |  |
| *D. santomea* STO.7 | Daniel Matute |  |
| *D. santomea* DBAT400.2 | Daniel Matute |  |
| *D. santomea* Thera13 | Daniel Matute |  |
| *D. yakuba* 4-26 | Daniel Matute |  |
| *D. yakuba* 1235.4 | Daniel Matute |  |
| *D. yakuba* NY62 | US Stock centre |  |
| *D. teissieri* TUZ11 | Daniel Matute |  |
| *D. teissieri* Selinda | Daniel Matute |  |
| *D. teissieri* CT03 | Daniel Matute |  |
| *D. melanogaster* w1118 |  |  |
| *D. melanogaster* Canton-S |  |  |
| *D. melanogaster* B12 | Global Diversity Line |  |
| **Assay materials** |  |  |
| Sucrose | Sigma-Aldrich | S5390 |
| Activated charcoal | Sigma-Aldrich | 242276 |
| AgaPure Agarose | Canvax | AG005 |
| Aluminium plate<br>0.5mm thickness | Alfer / Jumbo | 4.151.908 |
| Microcontroller board | Arduino | Uno Rev3 |
| Temperature sensors | <u>Variohm</u><br><u>EuroSensor</u> | ERTD-PT-1000-A-3850 |
| Peltier elements | Hebei | TEC1-12705 |
| Bench top power<br>supplies | PeakTech | P6080 |
| Camera | Teledyne Flir | CM3-U3-50S5M-CS |
| Lens | Computar | M1620-MPW2 |
| Thermal camera | Teledyne Flir | Lepton 3.5 |
| **Software and datasets** |  |  |
| Flye | <sup>84</sup> | https://github.com/fenderglass/Flye |
| Purge Haplotigs | <sup>85</sup> | https://bitbucket.org/mroachawri/purge_haplotigs |
| Pilon | <sup>86</sup> | https://github.com/broadinstitute/pilon |
| STAR | <sup>87</sup> | https://github.com/alexdobin/STAR |
| OrthoFinder | <sup>88</sup> | https://github.com/davidemms/OrthoFinder |
| RepeatModeler2.0 | <sup>89</sup> | https://github.com/Dfam-consortium/RepeatModeler |
| RepeatMasker | <sup>90</sup> | https://www.repeatmasker.org/RepeatMasker/ |
| BRAKER pipeline | <sup>91,92</sup> | https://github.com/Gaius-Augustus/BRAKER |
| MEGA11 | <sup>93</sup> | https://www.megasoftware.net/ |
| Arthropoda odb v10 | <sup>94</sup> | https://busco.ezlab.org/frames/art |
| FIMtrack | <sup>95</sup> | https://www.uni-muenster.de/Geoinformatics.cvmls/media/fim-media.html |
| Spinnaker SDK | Teledyne Flir | Spinnaker-SDK |
| **Python packages** |  |  |
| opencv2 | <sup>96</sup> | https://pypi.org/project/opencv-python/ |
| pandas | <sup>97,98</sup> | https://pypi.org/project/pandas/ |
| numpy | <sup>99</sup> | https://pypi.org/project/numpy/ |
| SciPy | <sup>100</sup> | https://pypi.org/project/scipy/ |
| Larvaworld | <sup>71</sup> | https://pypi.org/project/larvaworld/ |
| **R packages** |  |  |
| ape | <sup>101</sup> | https://cran.r-project.org/web/packages/ape/index.html |
| geiger | <sup>102</sup> | https://cran.r-project.org/web/packages/geiger/index.html |
| phylolm | <sup>103</sup> | https://cran.r-project.org/web/packages/phylolm/index.html |
| phytools | <sup>56</sup> | https://cran.r-project.org/web/packages/phytools/index.html |
| emmeans |  | https://cran.r-project.org/web/packages/emmeans/index.html |
| R 4.2.2 | <sup>104</sup> | https://www.r-project.org/ |

## Supplemental Information

**Document S1.** Figures S1–S7 and Tables S1, S3–S6

**Document S2.** Tables S2–S3

Table S2 Electivity measures for each strain used in study.

Table S3 Linear regression results for thermotactic behaviours.

## Acknowledgements

We thank Gwénaëlle Bontonou and Bastien Saint-Leandre for their insightful feedback throughout the course of this work and on earlier comments on the manuscript; Simon Sprecher, Tim-Henning Humberg, Martin Kapun, and Anton Strunov for valuable discussions on arena builds and larval tracking; the Swiss Institute of Bioinformatics for providing computational resources; Koichiro Tamura, Bruno Lemaitre, Benjamin Prud’homme, and Daniel Matute for generously providing fly specimens; and Tess Baticle and Afrah Hassan assisting in fly work and maintence. P.S. is funded through the ‘‘iBehave’’ research consortium (https://ibehave.nrw/) from the program ‘‘Netzwerke 2021’’ an initiative of the Ministry of Culture and Science of the State of North Rhine-Westphalia, Germany. Research in JRA’s laboratory was supported by the University of Lausanne and the Swiss National Science Foundation (grants PP00P3_176956 and 310030_201188).

## Author contributions

Conceptualisation: T.K. and J.R.A.; Methodology: T.K., P.S., and J.R.A.; Software: T.K. and P.S.; Formal Analysis: T.K. and J.R.A.; Investigation: T.K., M.G., and J.R.A.; Data Curation: T.K. and J.R.A.; Writing – Original Draft: T.K.; Writing – Reviewing and Editing: T.K., P.S., M.P.N., and J.R.A.; Funding Acquisition: J.R.A.

## References

1. Stratman, R., and Markow, T.A. (1998). Resistance to thermal stress in desert *Drosophila*: Desert Drosophila thermotolerance. Functional Ecology 12, 965–970. 10.1046/j.1365-2435.1998.00270.x.

2. Nagy, K.A. (1973). Behavior, Diet and Reproduction in a Desert Lizard, Sauromalus obesus. Copeia 1973, 93. 10.2307/1442363.

3. Schmidt-Nielsen, K., and Schmidt-Nielsen, B. (1952). Water metabolism of desert mammals. Physiological Reviews 32, 135–166.

4. De Jong, H., and Ciliberti, P. (2014). How cold-adapted flightless flies dispersed over the northern hemisphere: phylogeny and biogeography of the snow fly genus *Chionea* Dalman (Diptera: Limoniidae): Phylogeny and biogeography of the snow fly genus Chionea. Syst Entomol 39, 563–589. 10.1111/syen.12075.

5. Bista, I., Wood, J.M.D., Desvignes, T., McCarthy, S.A., Matschiner, M., Ning, Z., Tracey, A., Torrance, J., Sims, Y., Chow, W., et al. (2023). Genomics of cold adaptations in the Antarctic notothenioid fish radiation. Nat Commun 14, 3412. 10.1038/s41467-023-38567-6.

6. Pirri, F., Ometto, L., Fuselli, S., Fernandes, F.A.N., Ancona, L., Perta, N., Di Marino, D., Le Bohec, C., Zane, L., and Trucchi, E. (2022). Selection-driven adaptation to the extreme Antarctic environment in the Emperor penguin. Heredity 129, 317–326. 10.1038/s41437-022-00564-8.

7. Seebacher, F. (2009). Responses to temperature variation: integration of thermoregulation and metabolism in vertebrates. Journal of Experimental Biology 212, 2885–2891. 10.1242/jeb.024430.

8. Rezende, E.L., and Bacigalupe, L.D. (2015). Thermoregulation in endotherms: physiological principles and ecological consequences. J Comp Physiol B 185, 709–727. 10.1007/s00360-015-0909-5.

9. Kearney, M., Shine, R., and Porter, W.P. (2009). The potential for behavioral thermoregulation to buffer “cold-blooded” animals against climate warming. Proceedings of the National Academy of Sciences 106, 3835–3840. 10.1073/pnas.0808913106.

10. Heinrich, B. (1974). Thermoregulation in Endothermic Insects: Body temperature is closely attuned to activity and energy supplies. Science 185, 747–756. 10.1126/science.185.4153.747.

11. Stevenson, R.D. (1985). Body Size and Limits to the Daily Range of Body Temperature in Terrestrial Ectotherms. The American Naturalist 25, 102–117.

12. Garrity, P.A., Goodman, M.B., Samuel, A.D., and Sengupta, P. (2010). Running hot and cold: behavioral strategies, neural circuits, and the molecular machinery for thermotaxis in *C. elegans* and *Drosophila*. Genes Dev. 24, 2365–2382. 10.1101/gad.1953710.

13. Sømme, L. (1989). Adaptations of Terrestrial Arthropods to the Alpine Environment. Biological Reviews 64, 367–407. 10.1111/j.1469-185X.1989.tb00681.x.

14. Ward, D., and Seely, M.K. (1996). Behavioral Thermoregulation of Six Namib Desert Tenebrionid Beetle Species (Coleoptera). Annals of the Entomological Society of America 89, 442–451. 10.1093/aesa/89.3.442.

15. Ma, G., and Ma, C.-S. (2012). Effect of acclimation on heat-escape temperatures of two aphid species: Implications for estimating behavioral response of insects to climate warming. Journal of Insect Physiology 58, 303–309. 10.1016/j.jinsphys.2011.09.003.

16. Malmos, K.G., Lüdeking, A.H., Vosegaard, T., Aagaard, A., Bechsgaard, J., Sørensen, J.G., and Bilde, T. (2021). Behavioural and physiological responses to thermal stress in a social spider. Functional Ecology 35, 2728–2742. 10.1111/1365-2435.13921.

17. Harvey, J.A., Tougeron, K., Gols, R., Heinen, R., Abarca, M., Abram, P.K., Basset, Y., Berg, M., Boggs, C., Brodeur, J., et al. (2023). Scientists’ warning on climate change and insects. Ecological Monographs 93. 10.1002/ecm.1553.

18. Dobzhansky, T. (1943). Genetics of Natural Populations IX. Temporal Changes in the Composition of Populations of Drosophila pseudoobscura. Genetics 28, 162–186. 10.1093/genetics/28.2.162.

19. Dobzhansky, T. (1950). Genetics of Natural Populations. XIX. Origin of Heterosis through Natural Selection in Populations of Drosophila pseudoobscura. Genetics 35, 288–302. 10.1093/genetics/35.3.288.

20. Rego, C., Balanyà, J., Fragata, I., Matos, M., Rezende, E.L., and Santos, M. (2010). Clinal Patterns of Chromosomal Inversion Polymorphisms in Drosophila subobscura are Partly Associated with Thermal Preferences and Heat Stress Resistance. Evolution 64, 385–397. 10.1111/j.1558-5646.2009.00835.x.

21. Kapun, M., Fabian, D.K., Goudet, J., and Flatt, T. (2016). Genomic Evidence for Adaptive Inversion Clines in *Drosophila melanogaster*. Mol Biol Evol 33, 1317–1336. 10.1093/molbev/msw016.

22. Kapun, M., and Flatt, T. (2019). The adaptive significance of chromosomal inversion polymorphisms in *Drosophila melanogaster*. Mol Ecol 28, 1263–1282. 10.1111/mec.14871.

23. Fuller, Z.L., Koury, S.A., Phadnis, N., and Schaeffer, S.W. (2019). How chromosomal rearrangements shape adaptation and speciation: Case studies in *Drosophila pseudoobscura* and its sibling species *Drosophila persimilis*. Mol Ecol 28, 1283–1301. 10.1111/mec.14923.

24. Parsons, P.A. (1977). Genes, Behavior, and Evolutionary Processes: The Genus Drosophila. In Advances in Genetics, E. W. Caspari, ed. (Academic Press), pp. 1–32. 10.1016/S0065-2660(08)60244-8.

25. Gibert, P., Moreteau, B., Pétavy, G., Karan, D., and David, J.R. (2001). Chill-Coma Tolerance, a Major Climatic Adaptation Among Drosophila Species. Evol 55, 1063. http://doi.org/10.1554/0014-3820(2001)055[1063:CCTAMC]2.0.CO;2.

26. Kellerman, V., Overgaard, J., Fløjgaard, C., Svenning, J.-C., and Loeschcke, V. (2012). Upper thermal limits of Drosophila are linked to species distributions and strongly constrained phylogenetically. PNAS 109, 16228–16233.

27. Michalak, P., Minkov, I., Helin, A., Lerman, D.N., Bettencourt, B.R., Feder, M.E., Korol, A.B., and Nevo, E. (2001). Genetic evidence for adaptation-driven incipient speciation of Drosophila melanogaster along a microclimatic contrast in “Evolution Canyon,” Israel. Proceedings of the National Academy of Sciences 98, 13195–13200. 10.1073/pnas.231478298.

28. Ito, F., and Awasaki, T. (2022). Comparative analysis of temperature preference behavior and effects of temperature on daily behavior in 11 Drosophila species. Sci Rep 12, 12692. 10.1038/s41598-022-16897-7.

29. Soto-Yéber, L., Soto-Ortiz, J., Godoy, P., and Godoy-Herrera, R. (2018). The behavior of adult Drosophila in the wild. PLoS ONE 13, e0209917. 10.1371/journal.pone.0209917.

30. Kimura, M.T. (1988). Adaptations to Temperate Climates and Evolution of Overwintering Strategies in the Drosophila Melanogaster Species Group. Evolution 42, 1288–1297. 10.1111/j.1558-5646.1988.tb04188.x.

31. Kimura, M.T. (2004). Cold and heat tolerance of drosophilid flies with reference to their latitudinal distributions. Oecologia 140, 442–449. 10.1007/s00442-004-1605-4.

32. Goda, T., Leslie, J.R., and Hamada, F.N. (2014). Design and Analysis of Temperature Preference Behavior and its Circadian Rhythm in Drosophila. JoVE, 51097. 10.3791/51097.

33. Klein, M., Afonso, B., Vonner, A.J., Hernandez-Nunez, L., Berck, M., Tabone, C.J., Kane, E.A., Pieribone, V.A., Nitabach, M.N., Cardona, A., et al. (2015). Sensory determinants of behavioral dynamics in *Drosophila* thermotaxis. Proc. Natl. Acad. Sci. U.S.A. 112. 10.1073/pnas.1416212112.

34. Frank, D.D., Jouandet, G.C., Kearney, P.J., Macpherson, L.J., and Gallio, M. (2015). Temperature representation in the Drosophila brain. Nature 519, 358–361. 10.1038/nature14284.

35. Tratter Kinzner, M., Kinzner, M.-C., Kaufmann, R., Hoffmann, A.A., Arthofer, W., Schlick-Steiner, B.C., and Steiner, F.M. (2019). Is temperature preference in the laboratory ecologically relevant for the field? The case of Drosophila nigrosparsa. Global Ecology and Conservation 18, e00638. 10.1016/j.gecco.2019.e00638.

36. Abram, P.K., Boivin, G., Moiroux, J., and Brodeur, J. (2017). Behavioural effects of temperature on ectothermic animals: unifying thermal physiology and behavioural plasticity: Effects of temperature on animal behaviour. Biol Rev 92, 1859–1876. 10.1111/brv.12312.

37. Gallio, M., Ofstad, T.A., Macpherson, L.J., Wang, J.W., and Zuker, C.S. (2011). The Coding of Temperature in the Drosophila Brain. Cell 144, 614–624. 10.1016/j.cell.2011.01.028.

38. Budelli, G., Ni, L., Berciu, C., Van Giesen, L., Knecht, Z.A., Chang, E.C., Kaminski, B., Silbering, A.F., Samuel, A., Klein, M., et al. (2019). Ionotropic Receptors Specify the Morphogenesis of Phasic Sensors Controlling Rapid Thermal Preference in Drosophila. Neuron 101, 738–747.e3. 10.1016/j.neuron.2018.12.022.

39. Hamada, F.N., Rosenzweig, M., Kang, K., Pulver, S.R., Ghezzi, A., Jegla, T.J., and Garrity, P.A. (2008). An internal thermal sensor controlling temperature preference in Drosophila. Nature 454, 217–220. 10.1038/nature07001.

40. Hernandez-Nunez, L., Chen, A., Budelli, G., Berck, M.E., Richter, V., Rist, A., Thum, A.S., Cardona, A., Klein, M., Garrity, P., et al. (2021). Synchronous and opponent thermosensors use flexible cross-inhibition to orchestrate thermal homeostasis. Science Advances 7, eabg6707. 10.1126/sciadv.abg6707.

41. Liu, L., Yermolaieva, O., Johnson, W.A., Abboud, F.M., and Welsh, M.J. (2003). Identification and function of thermosensory neurons in Drosophila larvae. Nat Neurosci 6, 267–273. 10.1038/nn1009.

42. Hwang, R.Y., Zhong, L., Xu, Y., Johnson, T., Zhang, F., Deisseroth, K., and Tracey, W.D. (2007). Nociceptive Neurons Protect Drosophila Larvae from Parasitoid Wasps. Current Biology 17, 2105–2116. 10.1016/j.cub.2007.11.029.

43. Turner, H.N., Armengol, K., Patel, A.A., Himmel, N.J., Sullivan, L., Iyer, S.C., Bhattacharya, S., Iyer, E.P.R., Landry, C., Galko, M.J., et al. (2016). The TRP Channels Pkd2, NompC, and Trpm Act in Cold-Sensing Neurons to Mediate Unique Aversive Behaviors to Noxious Cold in Drosophila. Current Biology 26, 3116–3128. 10.1016/j.cub.2016.09.038.

44. Barbagallo, B., and Garrity, P.A. (2015). Temperature sensation in Drosophila. Current Opinion in Neurobiology 34, 8–13. 10.1016/j.conb.2015.01.002.

45. Xiao, R., and Xu, X.Z.S. (2021). Temperature Sensation: From Molecular Thermosensors to Neural Circuits and Coding Principles. Annu. Rev. Physiol. 83, 205–230. 10.1146/annurev-physiol-031220-095215.

46. Kimura, M.T. (1982). Cold Hardiness and Preimaginal Period in Two Closely Related Species, Drosophila takahashii and D. lutescens. Kontyu 50, 638–648.

47. Lachaise, D., Cariou, M.-L., David, J.R., Lemeunier, F., Tsacas, L., and Ashburner, M. (1988). Historical Biogeography of the Drosophila melanogaster Species Subgroup. In Evolutionary Biology, M. K. Hecht, B. Wallace, and G. T. Prance, eds. (Springer US), pp. 159–225. 10.1007/978-1-4613-0931-4_4.

48. Lachaise, D., Harry, M., Solignac, M., Lemeunier, F., Bénassi, V., and Cariou, M.-L. (2000). Evolutionary novelties in islands: *Drosophila santomea* , a new *melanogaster* sister species from São Tomé. Proc. R. Soc. Lond. B 267, 1487–1495. 10.1098/rspb.2000.1169.

49. MacLean, H.J., Overgaard, J., Kristensen, T.N., Lyster, C., Hessner, L., Olsvig, E., and Sørensen, J.G. (2019). Temperature preference across life stages and acclimation temperatures investigated in four species of Drosophila. Journal of Thermal Biology 86, 102428. 10.1016/j.jtherbio.2019.102428.

50. Huda, A., Omelchenko, A.A., Vaden, T.J., Castaneda, A.N., and Ni, L. (2022). Responses of different Drosophila species to temperature changes. Journal of Experimental Biology 225, jeb243708. 10.1242/jeb.243708.

51. Turissini, D.A., Liu, G., David, J.R., and Matute, D.R. (2015). The evolution of reproductive isolation in the Drosophila yakuba complex of species. J Evol Biol 28, 557–575. 10.1111/jeb.12588.

52. Matute, D.R., Novak, C.J., and Coyne, J.A. (2009). Temperature-based extrinsic reproductive isolation in two species of *Drosophila*. Evolution 63, 595–612. 10.1111/j.1558-5646.2008.00588.x.

53. Cooper, B.S., Sedghifar, A., Nash, W.T., Comeault, A.A., and Matute, D.R. (2018). A Maladaptive Combination of Traits Contributes to the Maintenance of a Drosophila Hybrid Zone. Current Biology 28, 2940–2947.e6. 10.1016/j.cub.2018.07.005.

54. Fukatami, A. (1984). Cold temperature resistance in Drosophila lutescens and D. takahashii. Jpn J Genet. 59, 61–70. 10.1266/jjg.59.61.

55. Luo, L., Gershow, M., Rosenzweig, M., Kang, K., Fang-Yen, C., Garrity, P.A., and Samuel, A.D.T. (2010). Navigational Decision Making in Drosophila Thermotaxis. J Neurosci 30, 4261– 4272. 10.1523/JNEUROSCI.4090-09.2010.

56. Pincebourde, S., Murdock, C.C., Vickers, M., and Sears, M.W. (2016). Fine-Scale Microclimatic Variation Can Shape the Responses of Organisms to Global Change in Both Natural and Urban Environments. Integr. Comp. Biol. 56, 45–61. 10.1093/icb/icw016.

57. Pincebourde, S., and Woods, H.A. (2012). Climate uncertainty on leaf surfaces: the biophysics of leaf microclimates and their consequences for leaf-dwelling organisms. Functional Ecology 26, 844–853. 10.1111/j.1365-2435.2012.02013.x.

58. Ivlev, V.S. (1961). Experimental Ecology of the Feeding of Fishes (Yale University Press).

59. Arguello, J.R., Laurent, S., and Clark, A.G. (2019). Demographic History of the Human Commensal *Drosophila melanogaster*. Genome Biology and Evolution 11, 844–854. 10.1093/gbe/evz022.

60. Ørsted, I.V., and Ørsted, M. (2019). Species distribution models of the Spotted Wing *Drosophila* ( *Drosophila suzukii* , Diptera: Drosophilidae) in its native and invasive range reveal an ecological niche shift. J Appl Ecol 56, 423–435. 10.1111/1365-2664.13285.

61. Watanabe, T.K., and Kawanishi, M. (1983). Stasipatric speciation in Drosophila. Jpn J Genet. 58, 269–274. 10.1266/jjg.58.269.

62. Comeault, A., and Matute, D. (2020). Temperature-dependent competitive outcomes between the fruit flies Drosophila santomea and D. yakuba. Version 1 (Dryad). 10.5061/DRYAD.BK3J9KD8T https://doi.org/10.5061/DRYAD.BK3J9KD8T.

63. Loveless, J., and Webb, B. (2018). A Neuromechanical Model of Larval Chemotaxis. Integrative and Comparative Biology. 10.1093/icb/icy094.

64. Wystrach, A., Lagogiannis, K., and Webb, B. (2016). Continuous lateral oscillations as a core mechanism for taxis in Drosophila larvae. eLife 5, e15504. 10.7554/eLife.15504.

65. Gershow, M., Berck, M., Mathew, D., Luo, L., Kane, E.A., Carlson, J.R., and Samuel, A.D.T. (2012). Controlling airborne cues to study small animal navigation. Nat Methods 9, 290–296. 10.1038/nmeth.1853.

66. Soto-Padilla, A., Ruijsink, R., Sibon, O.C.M., Van Rijn, H., and Billeter, J.-C. (2018). Thermosensory perception regulates speed of movement in response to temperature changes in *Drosophila melanogaster*. Journal of Experimental Biology, jeb.174151. 10.1242/jeb.174151.

67. Mazzoni, E.O., Desplan, C., and Blau, J. (2005). Circadian Pacemaker Neurons Transmit and Modulate Visual Information to Control a Rapid Behavioral Response. Neuron 45, 293–300. 10.1016/j.neuron.2004.12.038.

68. Komarov, N., and Sprecher, S.G. (2022). The chemosensory system of the *Drosophila* larva: an overview of current understanding. Fly 16, 1–12. 10.1080/19336934.2021.1953364.

69. Jovanic, T., Winding, M., Cardona, A., Truman, J.W., Gershow, M., and Zlatic, M. (2019). Neural Substrates of Drosophila Larval Anemotaxis. Current Biology 29, 554–566.e4. 10.1016/j.cub.2019.01.009.

70. Kane, E.A., Gershow, M., Afonso, B., Larderet, I., Klein, M., Carter, A.R., De Bivort, B.L., Sprecher, S.G., and Samuel, A.D.T. (2013). Sensorimotor structure of *Drosophila* larva phototaxis. Proc. Natl. Acad. Sci. U.S.A. 110. 10.1073/pnas.1215295110.

71. Gomez-Marin, A., and Louis, M. (2014). Multilevel control of run orientation in Drosophila larval chemotaxis. Front. Behav. Neurosci. 8. 10.3389/fnbeh.2014.00038.

72. Knecht, Z.A., Silbering, A.F., Ni, L., Klein, M., Budelli, G., Bell, R., Abuin, L., Ferrer, A.J., Samuel, A.D., Benton, R., et al. (2016). Distinct combinations of variant ionotropic glutamate receptors mediate thermosensation and hygrosensation in Drosophila. eLife 5, e17879. 10.7554/eLife.17879.

73. Ni, L., Klein, M., Svec, K.V., Budelli, G., Chang, E.C., Ferrer, A.J., Benton, R., Samuel, A.D., and Garrity, P.A. (2016). The Ionotropic Receptors IR21a and IR25a mediate cool sensing in Drosophila. eLife 5, e13254. 10.7554/eLife.13254.

74. Wong, P.H., Braun, A., Malagarriga, D., Moehlis, J., Moreno-Bote, R., Pouget, A., and Louis, M. (2023). Computational principles of adaptive multisensory combination in the *Drosophila* larva (bioRxiv) 10.1101/2023.05.04.539474.

75. Sakagiannis, P., Jürgensen, A.-M., and Nawrot, M.P. (2024). A behavioral architecture for realistic simulations of Drosophila larva locomotion and foraging. Preprint at bioRxiv, 10.1101/2021.07.07.451470 https://doi.org/10.1101/2021.07.07.451470.

76. Malhi, Y., Franklin, J., Seddon, N., Solan, M., Turner, M.G., Field, C.B., and Knowlton, N. (2020). Climate change and ecosystems: threats, opportunities and solutions. Phil. Trans. R. Soc. B 375, 20190104. 10.1098/rstb.2019.0104.

77. Arguello, J.R., and Benton, R. (2017). Open questions: Tackling Darwin’s “instincts”: the genetic basis of behavioral evolution. BMC Biology 15. 10.1186/s12915-017-0369-3.

78. Pool, J.E., Corbett-Detig, R.B., Sugino, R.P., Stevens, K.A., Cardeno, C.M., Crepeau, M.W., Duchen, P., Emerson, J.J., Saelao, P., Begun, D.J., et al. (2012). Population Genomics of Sub-Saharan Drosophila melanogaster: African Diversity and Non-African Admixture. PLoS Genet 8, e1003080. 10.1371/journal.pgen.1003080.

79. Grenier, J.K., Arguello, J.R., Moreira, M.C., Gottipati, S., Mohammed, J., Hackett, S.R., Boughton, R., Greenberg, A.J., and Clark, A.G. (2015). Global Diversity Lines–A Five-Continent Reference Panel of Sequenced *Drosophila melanogaster* Strains. G3 Genes|Genomes|Genetics 5, 593–603. 10.1534/g3.114.015883.

80. Reilly, P.F. (2020). Population genomics of the D. yakuba clade. In Work, wealth, and well-being: Essays in macroeconomics, P. Andolfatto, ed. (Princeton University), pp. 62–121.

81. Lewald, K.M., Abrieux, A., Wilson, D.A., Lee, Y., Conner, W.R., Andreazza, F., Beers, E.H., Burrack, H.J., Daane, K.M., Diepenbrock, L., et al. (2021). Population genomics of *Drosophila suzukii* reveal longitudinal population structure and signals of migrations in and out of the continental United States. G3 Genes|Genomes|Genetics 11, jkab343. 10.1093/g3journal/jkab343.

82. Sokabe, T., Chen, H.-C., Luo, J., and Montell, C. (2016). A Switch in Thermal Preference in Drosophila Larvae Depends on Multiple Rhodopsins. Cell Reports 17, 336–344. 10.1016/j.celrep.2016.09.028.

83. Bieńkowski, A.O., and Orlova-Bienkowskaja, M.J. (2020). Invasive Agricultural Pest Drosophila suzukii (Diptera, Drosophilidae) Appeared in the Russian Caucasus. Insects 11, 826. 10.3390/insects11110826.

84. Estay, S.A., Silva, C.P., López, D.N., and Labra, F.A. (2023). Disentangling the spread dynamics of insect invasions using spatial networks. Front. Ecol. Evol. 11, 1124890. 10.3389/fevo.2023.1124890.

85. Hamby, K.A., E. Bellamy, D., Chiu, J.C., Lee, J.C., Walton, V.M., Wiman, N.G., York, R.M., and Biondi, A. (2016). Biotic and abiotic factors impacting development, behavior, phenology, and reproductive biology of Drosophila suzukii. J Pest Sci 89, 605–619. 10.1007/s10340-016-0756-5.

86. Winkler, A., Jung, J., Kleinhenz, B., and Racca, P. (2020). A review on temperature and humidity effects on *Drosophila suzukii* population dynamics. Agr Forest Entomol 22, 179–192. 10.1111/afe.12381.

87. Tyrrell, J.J., Wilbourne, J.T., Omelchenko, A.A., Yoon, J., and Ni, L. (2021). Ionotropic Receptor-dependent cool cells control the transition of temperature preference in Drosophila larvae. PLOS Genetics 17, e1009499. 10.1371/journal.pgen.1009499.

88. Kolmogorov, M., Yuan, J., Lin, Y., and Pevzner, P.A. (2019). Assembly of long, error-prone reads using repeat graphs. Nat Biotechnol 37, 540–546. 10.1038/s41587-019-0072-8.

89. Roach, M.J., Schmidt, S.A., and Borneman, A.R. (2018). Purge Haplotigs: allelic contig reassignment for third-gen diploid genome assemblies. BMC Bioinformatics 19, 460. 10.1186/s12859-018-2485-7.

90. Bontonou, G., Saint-Leandre, B., Kafle, T., Baticle, T., Hassan, A., Sánchez-Alcañiz, J.A., and Arguello, J.R. (2024). Evolution of chemosensory tissues and cells across ecologically diverse Drosophilids. Nat Commun 15, 1047. 10.1038/s41467-023-44558-4.

91. Dobin, A., Davis, C.A., Schlesinger, F., Drenkow, J., Zaleski, C., Jha, S., Batut, P., Chaisson, M., and Gingeras, T.R. (2013). STAR: ultrafast universal RNA-seq aligner. Bioinformatics 29, 15–21. 10.1093/bioinformatics/bts635.

92. Emms, D.M., and Kelly, S. (2019). OrthoFinder: phylogenetic orthology inference for comparative genomics. Genome Biol 20, 238. 10.1186/s13059-019-1832-y.

93. Flynn, J.M., Hubley, R., Goubert, C., Rosen, J., Clark, A.G., Feschotte, C., and Smit, A.F. (2020). RepeatModeler2 for automated genomic discovery of transposable element families. Proc. Natl. Acad. Sci. U.S.A. 117, 9451–9457. 10.1073/pnas.1921046117.

94. Smit, A.F., Hubley, R., and Green, P. (2013). RepeatMasker Open-4.0.

95. Hoff, K.J., Lange, S., Lomsadze, A., Borodovsky, M., and Stanke, M. (2016). BRAKER1: Unsupervised RNA-Seq-Based Genome Annotation with GeneMark-ET and AUGUSTUS. Bioinformatics 32, 767–769. 10.1093/bioinformatics/btv661.

96. Hoff, K.J., Lomsadze, A., Borodovsky, M., and Stanke, M. (2019). Whole-Genome Annotation with BRAKER. In Gene Prediction Methods in Molecular Biology., M. Kollmar, ed. (Springer New York), pp. 65–95. 10.1007/978-1-4939-9173-0_5.

97. Kriventseva, E.V., Kuznetsov, D., Tegenfeldt, F., Manni, M., Dias, R., Simão, F.A., and Zdobnov, E.M. (2019). OrthoDB v10: sampling the diversity of animal, plant, fungal, protist, bacterial and viral genomes for evolutionary and functional annotations of orthologs. Nucleic Acids Research 47, D807–D811. 10.1093/nar/gky1053.

98. Tamura, K., Stecher, G., and Kumar, S. (2021). MEGA11: Molecular Evolutionary Genetics Analysis Version 11. Molecular Biology and Evolution 38, 3022–3027. 10.1093/molbev/msab120.

99. Suvorov, A., Kim, B.Y., Wang, J., Armstrong, E.E., Peede, D., D’Agostino, E.R.R., Price, D.K., Waddell, P.J., Lang, M., Courtier-Orgogozo, V., et al. (2022). Widespread introgression across a phylogeny of 155 Drosophila genomes. Current Biology 32, 111–123.e5. 10.1016/j.cub.2021.10.052.

100. Warrick, J.M., Vakil, M.F., and Tompkins, L. (1999). Spectral Sensitivity of Wild-Type and Mutant *Drosophila Melanogaster* Larvae. Journal of Neurogenetics 13, 145–156. 10.3109/01677069909083471.

101. Xiang, Y., Yuan, Q., Vogt, N., Looger, L.L., Jan, L.Y., and Jan, Y.N. (2010). Light-avoidance-mediating photoreceptors tile the Drosophila larval body wall. Nature 468, 921–926. 10.1038/nature09576.

102. Risse, B., Berh, D., Otto, N., Klämbt, C., and Jiang, X. (2017). FIMTrack: An open source tracking and locomotion analysis software for small animals. PLoS Comput Biol 13, e1005530. 10.1371/journal.pcbi.1005530.

103. Sakagiannis, P., Jürgensen, A.-M., and Nawrot, M.P. (2021). A realistic locomotory model of *Drosophila* larva for behavioral simulations (bioRxiv) 10.1101/2021.07.07.451470.

104. Kandel E.R., Schwartz J.H., Jessell T.M., Siegelbaum S.A., Hudspeth A.J., and Mack S Principles of Neural Science, Fifth Edition | AccessBiomedical Science. McGraw Hill Medical. https://neurology.mhmedical.com/content.aspx?sectionid=59138139&bookid=1049.

