## supplemental figures 1 - 7 for "Evolution of temperature preference behaviour among drosophilids"

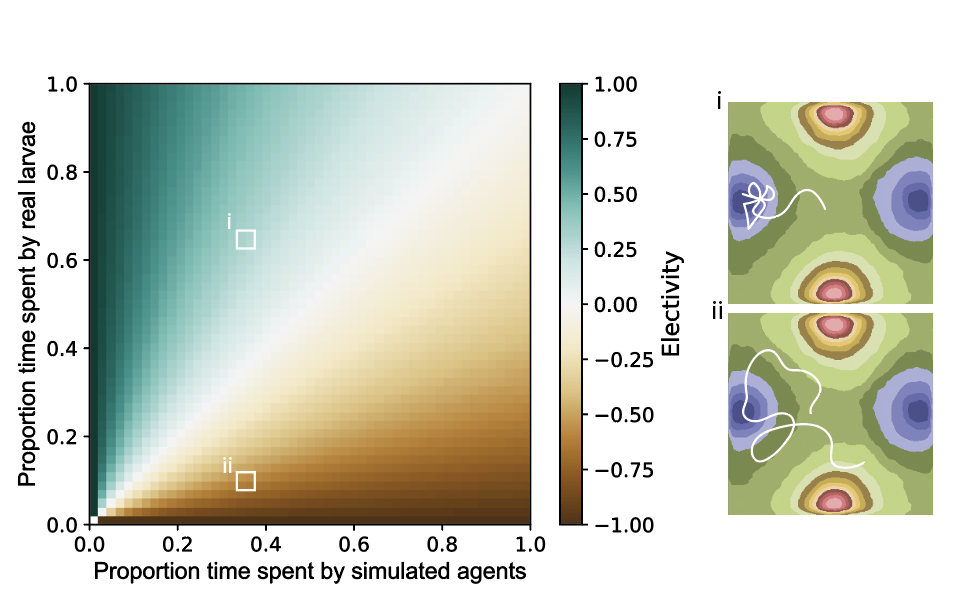

**Figure S1** Influence of temperature bins on electivity values. The left panel is adapted and modernised from Lechowicz (1982). It illustrates the impact of temperature distribution within the arena, that affects both the proportion time spent by simulated agents and larvae on the arena, which ultimately impacts the final electivity values. As bins are unequal in size, randomly moving simulated agents will spend more time in larger bins, and this effects the electivity values possible for those temperature bins when accounting for the time larvae spend too. Also, as larvae are always not present in n-1 bins, values tend to be towards the lower numbers. Consequently, electivity values tend to be closer to zero and are often skewed towards negative values. Panels on the right represent example larval tracks, with the corresponding points on the left panel, to show how larval movement can affect electivity values, in this case, for the blue zone.

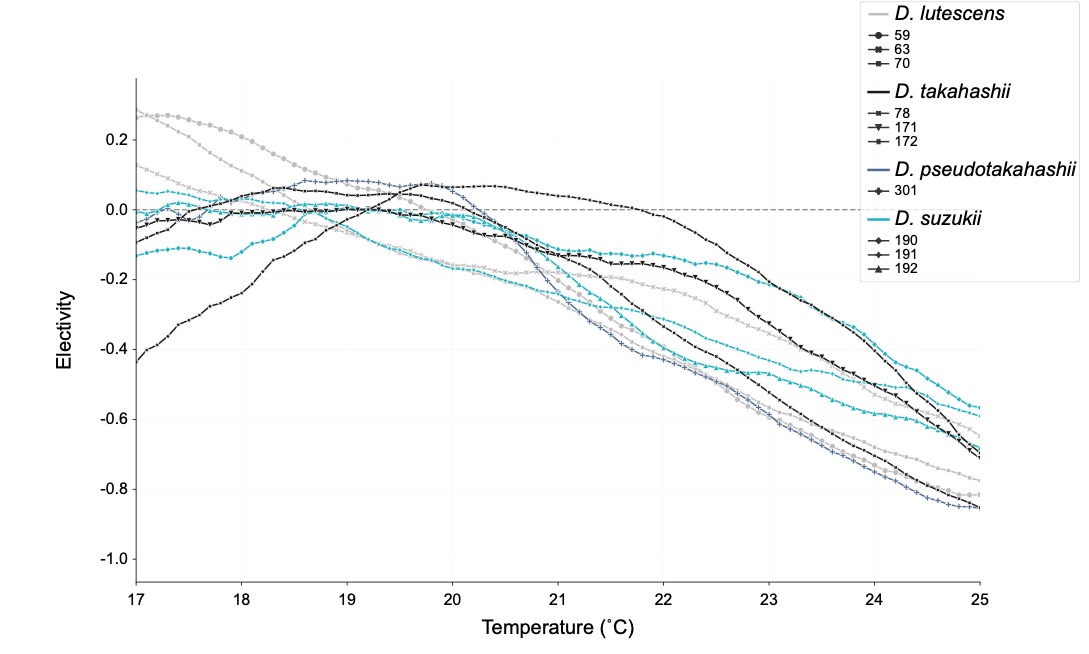

**Figure S2**
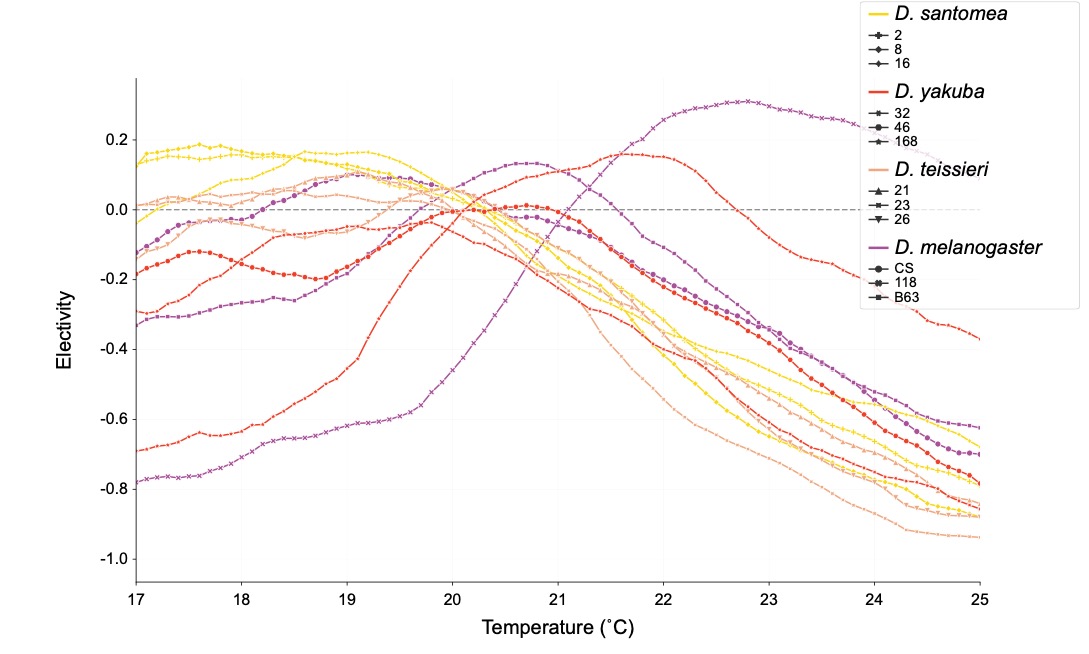

**Figure S3**

**Figure S2 and S3**: Electivity patterns across temperature for different strains. These lineplots illustrate the electivity patterns of strains used for each species across a range of temperatures. While there is variation among strains within a species, they often exhibit similar electivity patterns. Notably *D. melanogaster* w1118, *D. yakuba* 168, and *D. takahashii* 172 deviate from the common profile amongst the other two strains in the species.

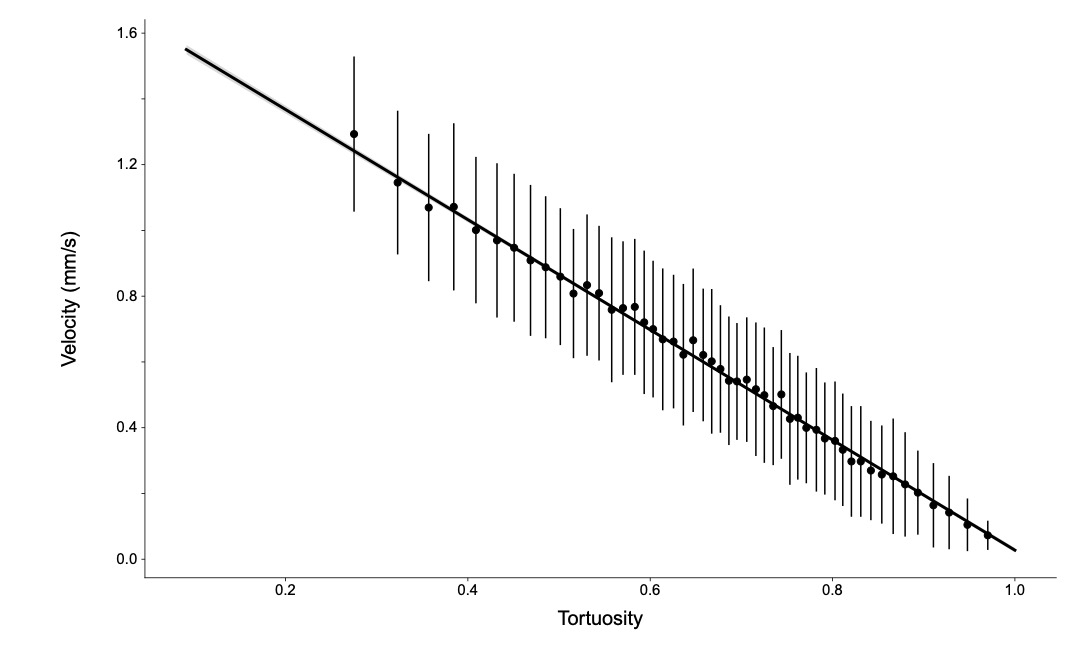

**Figure S4** Correlation between velocity and tortuosity. This figure ullustrates the relationship between velocity and tortuosity across all assays, demonstrating a clear correlation. It shows that a less tortuous, or straighter, run is associated with faster movemement. This correlation suggests that the larval movement in terms of tortuosity directly influences the speed at which a larva moves.

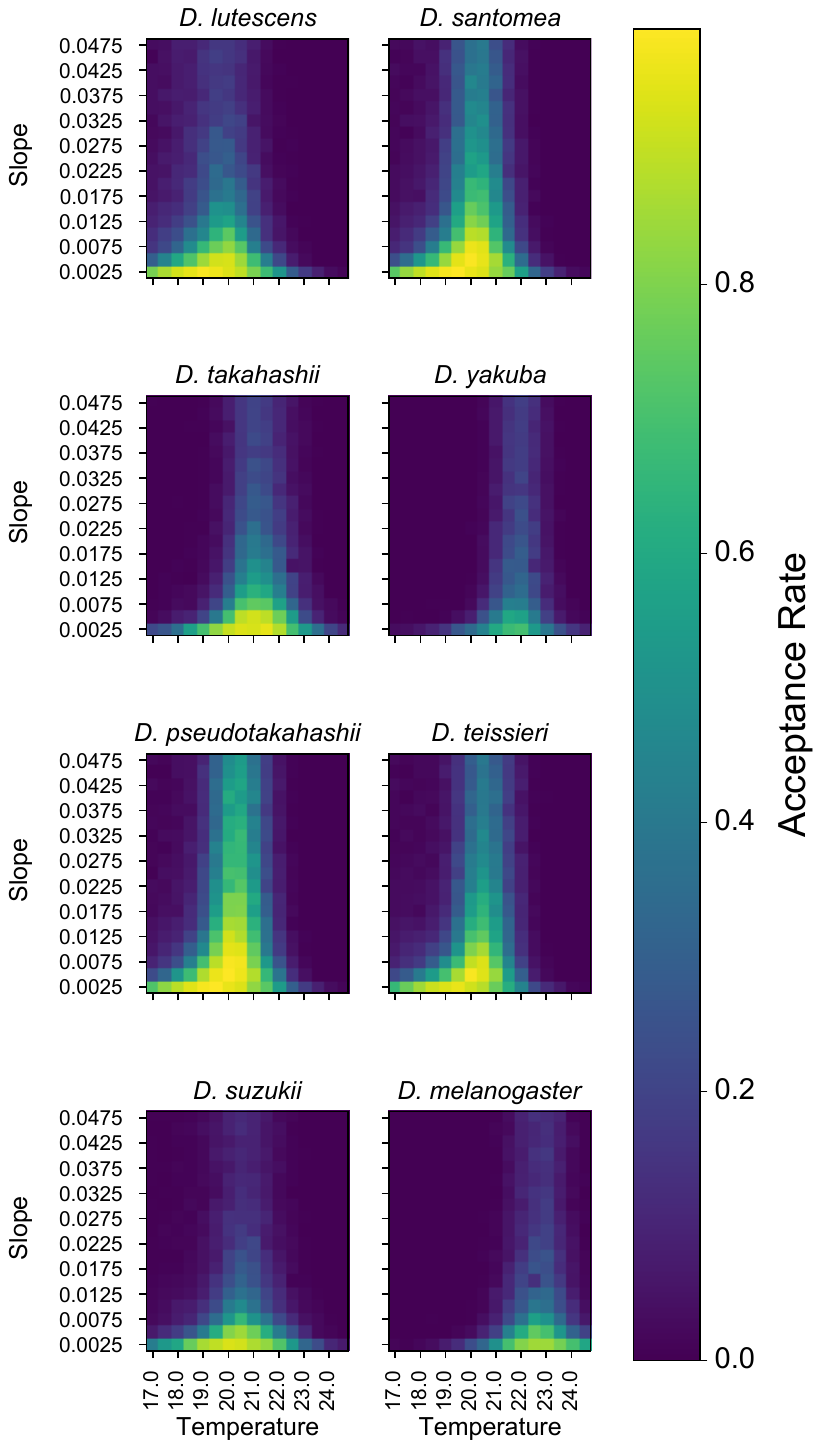

**Figure S5** Preliminary grid search for optimal slope values. In this preliminary grid search, the aim was to identify the most suitable slope values from a broad range of slope values, to focus in on the parameter values for the final simulations. The results suggest that lower slope values were optimal for all species in this grid search. This indicates a broader preference for temperatures around the preferred temperature within this innocuous range.

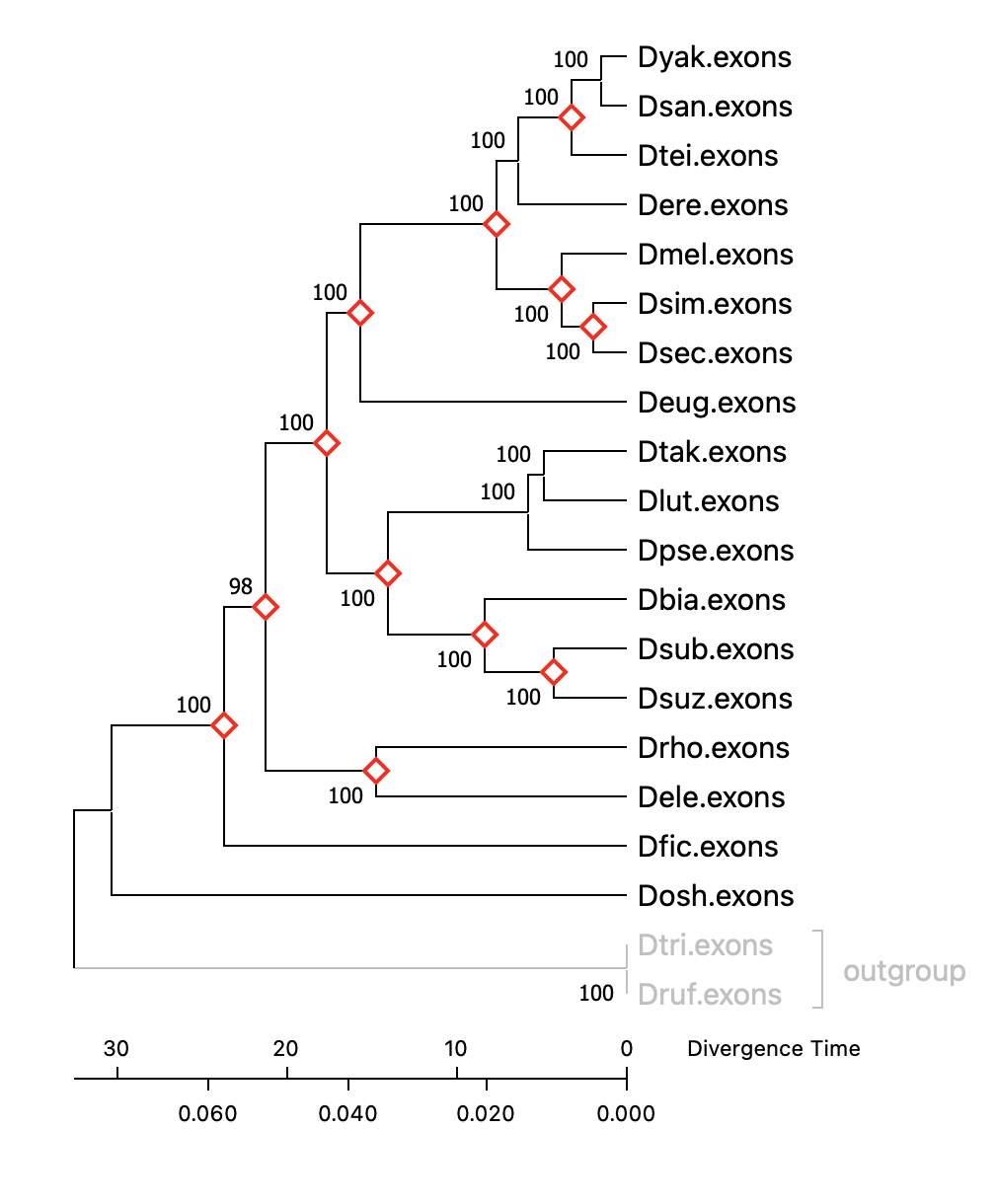

**Figure S6** Dated phylogeny generated in MEGA11. Red diamonds indicate dates that were input as calibration points from Table S6.

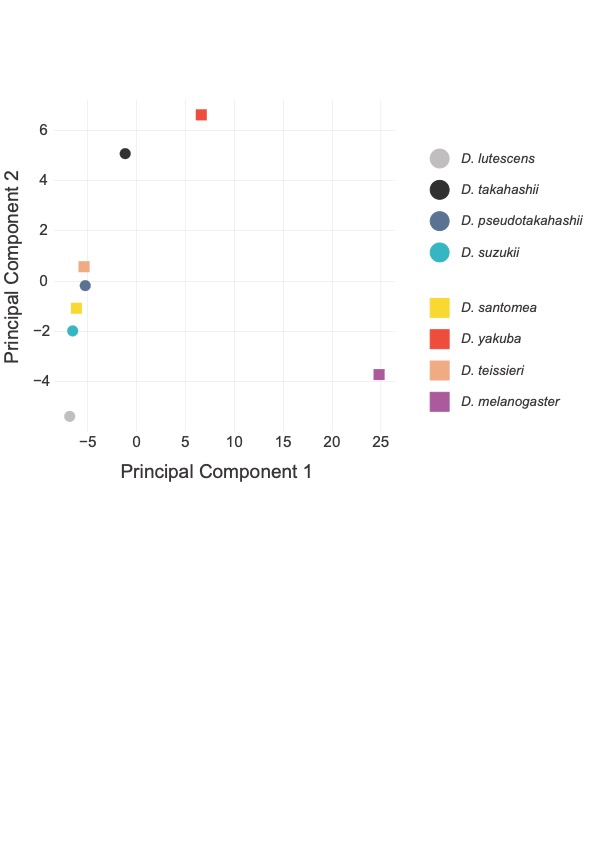

**Figure S7** PCA plot generated using grid values of simulations in Fig3A-H. The first axis separates *D. melanogaster* from all other species, with it having a homeostatic set point at higher temperatures than all other species. *Drosophila suzukii, D. pseudotakahashii*, *D. santomea* and *D. teissieri* sit very closely on both principal component axes, indicating that they have very similar fits to parameter sets.
